# TIGAR DEFICIENCY ENHANCES CARDIAC RESILIENCE THROUGH EPIGENETIC PROGRAMMING OF PARKIN EXPRESSION

**DOI:** 10.1101/2025.07.14.664830

**Authors:** Yan Tang, Stanislovas S. Jankauskas, Li Liu, Xujun Wang, Alus Xiaoli, Fajun Yang, Gaetano Santulli, Jeffrey E. Pessin

## Abstract

**BACKGROUND:** While mitochondrial dysfunction clearly drives cardiac deterioration in major heart diseases, the mechanisms controlling mitochondrial quality remain incompletely understood. This study investigated whether TIGAR (TP53-induced glycolysis and apoptosis regulator) deficiency influences cardiac protection through mitochondrial quality control pathways.

**METHODS:** We generated both whole-body and cardiomyocyte-specific TIGAR knockout mice that were assessed for cardiac function following myocardial infarction (induced by left anterior descending coronary artery ligation) and diet-induced cardiomyopathy (using a 6-month high-fat diet protocol). Mitochondrial quality control was evaluated through mitophagy assays, subcellular fractionation, and molecular analyses. Epigenetic regulation was assessed using whole-genome bisulfite sequencing, chromatin immunoprecipitation, and CRISPR-mediated gene editing in multiple cell lines.

**RESULTS:** Both whole-body (TKO) and cardiomyocyte-specific (hTKO) TIGAR knockout mice demonstrated cardioprotection following myocardial infarction. These animals maintained significantly better ejection fraction (43.35±17.76% vs 26.36±9.83% in wild-type controls, P<0.05) and displayed complete resistance to diet-induced cardiac hypertrophy, despite comparable weight gain. TIGAR deficiency led to dramatic increases in Parkin expression across multiple tissues, 6-fold increases in heart and muscle, and 5-fold increases in brain, which enhanced mitophagic responses during metabolic stress conditions including fasting and high-fat diet feeding. Generation of Parkin/TIGAR double knockout mice eliminated this protection, confirming Parkin’s essential role. Notably, adult manipulation of TIGAR through viral overexpression or knockdown failed to alter Parkin levels, suggesting developmental programming. Whole-genome bisulfite sequencing revealed reduced DNA methylation in a specific 3.2 kb region within Parkin gene intron 10, and CRISPR deletion of this regulatory region increased Parkin expression 10-fold in C2C12 myoblasts and 6-fold in 3T3-L1 fibroblasts.

**CONCLUSIONS:** These findings reveal a novel TIGAR-Parkin regulatory axis operating through epigenetic mechanisms during cardiac development to establish lifelong cardioprotection via enhanced mitochondrial quality control. This discovery points toward new therapeutic approaches targeting developmental metabolic programming for heart disease prevention and identifies specific epigenetic targets for cardiovascular therapy.

**CLINICAL PERSPECTIVE:** *What Is New?:*

- TIGAR deficiency establishes lifelong cardiac protection through developmental epigenetic programming of Parkin expression.
- A novel 3.2 kb differentially methylated region in Parkin intron 10 regulates mitochondrial quality control in the heart.
- Early metabolic programming during cardiac development can establish permanent cardioprotective phenotypes.
- The TIGAR-Parkin axis provides protection against both acute ischemic injury and chronic metabolic cardiomyopathy.

*What Are the Clinical Implications?:*

- Targeting the TIGAR-Parkin pathway could provide novel therapeutic approaches for preventing both ischemic heart disease and diabetic cardiomyopathy.
- Early developmental interventions targeting cardiac metabolism might establish lifelong cardiovascular protection.
- Epigenetic modifications of mitochondrial quality control genes represent potential therapeutic targets.
- The findings suggest optimal timing for cardiovascular preventive interventions may be during critical developmental windows.

## INTRODUCTION

Cardiovascular disease remains the leading cause of death worldwide, with mitochondrial dysfunction at the center of major cardiac pathologies.^1–4^ The heart, as one of the most metabolically active organs, requires exceptional mitochondrial function to meet its enormous energy demands, consuming approximately 6 kg of ATP daily.^5^ Impaired mitochondrial quality control mechanisms have serious consequences for cardiac performance and stress survival.

### Mitochondrial Dysfunction in Cardiac Disease

In ischemic heart disease, mitochondrial dysfunction drives cardiac pathology through interconnected mechanisms. During myocardial infarction, mitochondrial damage leads to accumulation of dysfunctional organelles, increased reactive oxygen species production, and release of mitochondrial damage-associated molecular patterns (DAMPs) that trigger robust inflammatory responses.^6, 7^ These deleterious processes create a vicious cycle where mitochondrial damage propagates cellular dysfunction beyond the original injury into viable myocardium, contributing to adverse remodeling and eventual heart failure.

The complexity is equally evident in diabetic heart disease, where mitochondrial dysfunction precedes overt cardiac functional abnormalities.^4, 8, 9^ Even with optimal glycemic control, diabetic patients remain at increased heart failure risk due to fundamental alterations in cardiac energy metabolism, including preferential fatty acid over glucose utilization, impaired energetic efficiency, and progressive mitochondrial dysfunction. This metabolic inflexibility renders diabetic hearts particularly vulnerable to additional stressors and limits adaptive capacity during increased demands.

### Mitochondrial Quality Control and the PINK1/Parkin Pathway

Mitochondrial quality control mechanisms, particularly selective autophagy of damaged mitochondria (mitophagy), represent critical cardioprotective pathways.^4^ The PINK1/Parkin pathway constitutes the primary mechanism governing mitophagy in mammalian cells.^10^ Under normal conditions, PINK1 (PTEN-induced kinase 1) is imported into healthy mitochondria and rapidly degraded. However, when mitochondrial membrane potential is compromised, PINK1 accumulates on the outer mitochondrial membrane, where it phosphorylates ubiquitin and recruits the E3 ubiquitin ligase Parkin.^11^ Activated Parkin then ubiquitinates numerous outer mitochondrial membrane proteins, marking damaged organelles for selective autophagic degradation.

The cardiac significance of this pathway has been demonstrated across multiple experimental models and disease states. Parkin-deficient mice exhibit exacerbated cardiac dysfunction following myocardial infarction, with increased infarct size, impaired functional recovery, and accelerated progression to heart failure.^12, 13^ Conversely, cardiac-specific Parkin overexpression or pharmacological enhancement of Parkin activity provides substantial cardioprotection, preserving cardiac function and improving survival following ischemic injury.^14–16^

### Developmental Programming of Cardiac Metabolism

Postnatal cardiac development involves dramatic metabolic reprogramming from glycolysis-dependent fetal metabolism to fatty acid oxidation-dependent adult metabolism. This transition requires extensive mitochondrial biogenesis, maturation of respiratory complexes, and establishment of robust quality control mechanisms. Parkin plays a crucial role in this developmental process, with Parkin-mediated mitophagy directing perinatal cardiac metabolic maturation.^17^ Disruption of this developmental programming through genetic deletion of key mitochondrial regulators including Parkin, TFAM, or PGC-1α results in severe cardiac dysfunction and often perinatal lethality.^18, 19^

The concept of developmental programming suggests that early environmental influences can establish persistent phenotypic changes affecting disease susceptibility throughout life.^20, 21^ In the cardiovascular system, evidence indicates that metabolic conditions during critical developmental windows can permanently alter cardiac structure, function, and stress resistance.^22, 23^ However, the molecular mechanisms underlying such programming, particularly those involving mitochondrial quality control, remain incompletely understood.

### TIGAR and Cardiac Metabolism

TIGAR (TP53-induced glycolysis and apoptosis regulator) functions as a multifaceted regulator of cellular metabolism and stress responses. Originally identified as a p53 target gene,^24^ TIGAR exhibits both phosphatase activity toward fructose-2,6-bisphosphate and additional regulatory functions influencing cellular energetics, redox homeostasis, and stress resistance.^25^ In cardiac pathophysiology, TIGAR’s role appears context-dependent. The p53-TIGAR axis can exacerbate cardiac damage after myocardial infarction,^26^ while TIGAR ablation preserves myocardial energetics in pressure overload heart failure and reduces pathological cardiac hypertrophy.^27, 28^

Our previous investigations revealed that TIGAR deficiency enhances TCA cycle flux, increases mitochondrial respiratory capacity, and exhibits anti-inflammatory properties. ^29, 30^ These metabolic effects suggested potential connections to mitochondrial quality control mechanisms, prompting us to investigate relationships between TIGAR and Parkin-mediated mitophagy in cardiac protection.

In this study, we demonstrate that TIGAR knockout mice exhibit dramatically increased Parkin expression through developmental epigenetic programming, conferring significant cardioprotection against both acute ischemic injury and chronic metabolic stress. Using Parkin/TIGAR double knockout models, we confirmed that these protective effects depend on Parkin. Whole-genome bisulfite sequencing revealed specific differentially methylated regions in the Parkin gene body that regulate its expression.

## METHODS

Detailed Methods and Materials are presented in the Supplemental Material.

### Experimental Animals and Study Design

All procedures were approved by the Einstein Institutional Animal Care and Use Committee. Whole-body TIGAR knockout (TKO), cardiomyocyte-specific TIGAR knockout (hTKO), Parkin knockout (PKO), and Parkin/TIGAR double knockout (PTKO) mice were generated using established protocols. Mice were housed under standard conditions with 12-hour light/dark cycles.

### Cardiac Disease Models and Functional Assessment

Myocardial infarction was induced by permanent left anterior descending coronary artery ligation in 16-week-old male mice. Diet-induced cardiomyopathy was modeled using 6-month high-fat diet (60% calories from fat). Cardiac function was assessed by transthoracic echocardiography using a 12-MHz probe (Vevo 2100, VisualSonics) at baseline and specified intervals post-intervention.

### Molecular and Biochemical Analyses

Total RNA was extracted using Direct-zol RNA MiniPrep Plus kit with quantitative RT-PCR performed using TaqMan assays. Protein analysis was conducted using standard Western blotting with appropriate antibodies. Mitochondrial and cytosolic fractions were isolated by differential centrifugation. RNA sequencing was performed by independent service providers using Illumina platforms with bioinformatic analysis using standard pipelines.

### Epigenetic and Genomic Studies

Whole-genome bisulfite sequencing (WGBS) was performed on cardiac and testicular tissues to identify differentially methylated regions. Chromatin immunoprecipitation (ChIP) was conducted using anti-RNA polymerase II antibody to assess transcriptional activity at the Parkin promoter. Methylation data were analyzed using Bismark with visualization in Integrative Genomics Viewer.

### Cell Culture and Gene Editing

C2C12 myoblasts and 3T3-L1 fibroblasts were cultured under standard conditions. CRISPR/Cas9-mediated deletion of the identified regulatory region was performed using dual guide RNAs targeting the differentially methylated region in Parkin intron 10. Successful deletions were confirmed by PCR and functional effects assessed by quantitative RT-PCR.

### Statistical Analysis

Sample sizes were determined by power calculations based on preliminary data. Data are presented as mean ± standard deviation. Statistical comparisons used unpaired t-tests, Mann-Whitney U tests, or ANOVA with appropriate post-hoc corrections. Significance was defined as *p<0.05, **p<0.01, ***p<0.001, ****p<0.0001. Analyses were performed using GraphPad Prism software.

## RESULTS

### TIGAR Deficiency Confers Comprehensive Cardioprotection Against Myocardial Infarction

TIGAR-deficient mice demonstrated comprehensive cardioprotection against ischemic injury. Analysis of both whole-body TIGAR knockout (TKO) and cardiomyocyte-specific TIGAR knockout (hTKO) mice distinguished between cardiomyocyte-intrinsic effects versus systemic metabolic changes (Figure 1 and Figure S1).

**Figure 1.**
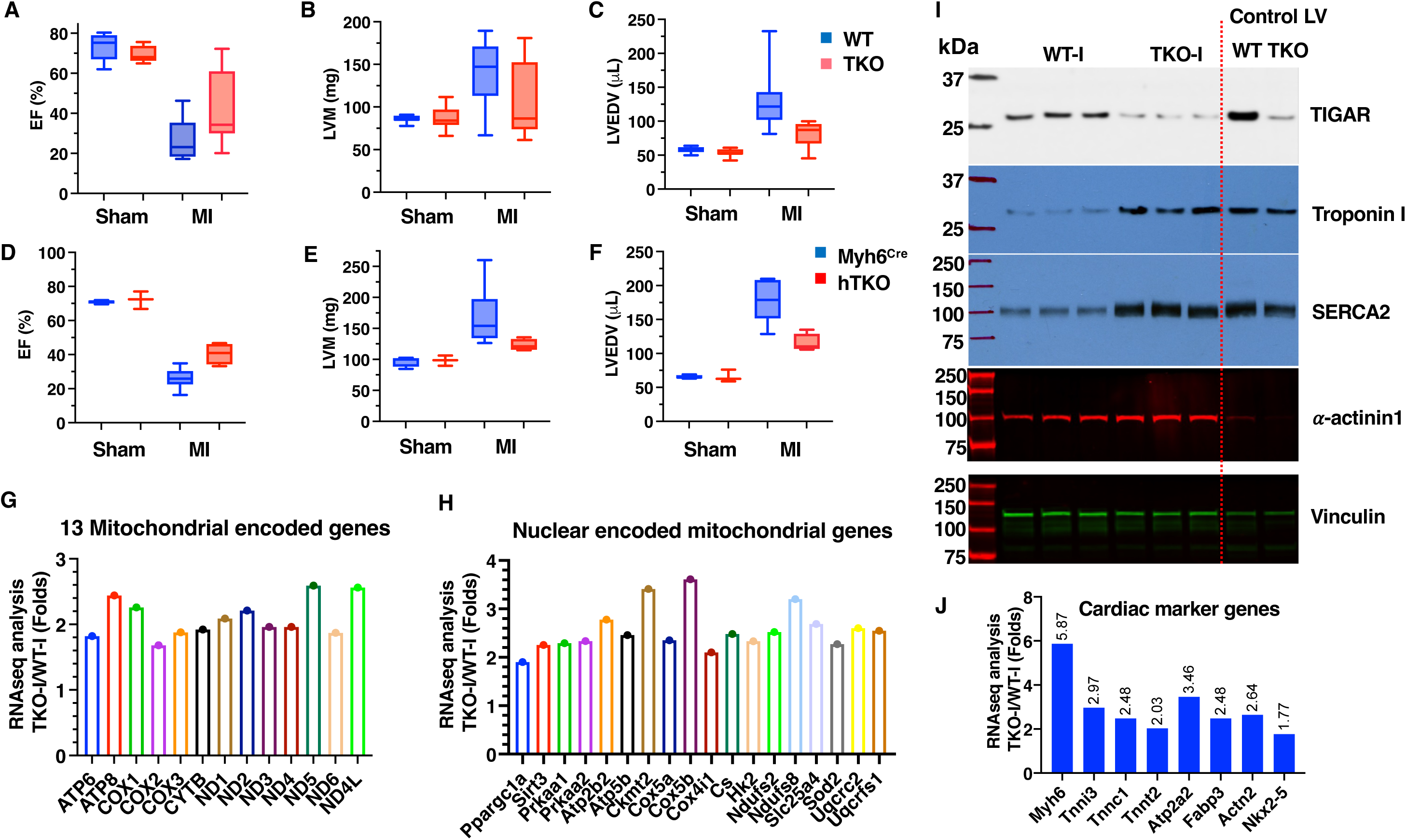
TIGAR knockout protects against post-myocardial infarction cardiac dysfunction. A-C, Echocardiograms and quantification of left ventricular ejection fraction (EF), mass (LVM), and end-diastolic volume (LVEDV) in 4-month-old male WT (n=10) and TKO mice (n=13) 4 weeks after myocardial infarction (MI). MI was induced by left anterior descending (LAD) artery ligation. Sham-operated mice served as controls. D-F, Echocardiograms and quantification of left ventricular ejection fraction (EF), mass (LVM), and end-diastolic volume (LVEDV) in 4-month-old male control Myh6Cre mice (n=6) and hTKO mice (n=4) 4 weeks after MI. G-H, RNA-seq analysis of mitochondrial-encoded genes (G) and nuclear-encoded mitochondrial genes (H) from the infarct zone, expressed as fold changes in TKO vs WT. I, Immunoblot analysis of TIGAR, Troponin I, SERCA2, α-actinin1, and Vinculin in infarcted heart tissue from WT and TKO mice, with normal left ventricular tissue (Control LV) as reference. J, RNA-seq analysis of cardiac marker genes from the infarct zone, expressed as fold changes in TKO vs WT. For RNA-seq analyses (G, H, J), RNA from 3 mice per group was pooled for sequencing. Data represent mean±SD; *P<0.05.

#### Preserved Cardiac Function Following Infarction

Following permanent left anterior descending coronary artery occlusion, both knockout models showed remarkably preserved cardiac function at 4 weeks post-injury. Our echocardiographic analysis revealed substantial preservation of cardiac function. TKO mice-maintained ejection fraction at 43.35±17.76% (n=13) while wild-type controls exhibited severely impaired function at 26.36±9.83% (n=10, P<0.05) (Figure 1A). This level of protection was substantial, representing retention of approximately 60% of baseline cardiac function compared to only 36% in controls.

Cardiomyocyte-specific knockout mice (hTKO) demonstrated similar protection with ejection fraction at 40.46±6.29% compared to control mice at 26.06±6.08% (Figure 1D). These findings indicated that cardioprotection resulted primarily from cardiomyocyte-intrinsic mechanisms rather than systemic metabolic alterations.

#### Reduced Pathological Remodeling

Both knockout models were significantly protected against increases in left ventricular mass and volume that typically follow myocardial infarction—key indicators of harmful cardiac remodeling (Figure 1B,C,E,F). While wild-type mice showed typical post-infarction changes including chamber dilation and compensatory wall thickening, knockout animals maintained relatively normal cardiac structure despite similar-sized infarct areas. This functional preservation was accompanied by significantly reduced cardiac enlargement in TKO hearts post-MI compared with wild-type hearts (Figure S2A,B). After MI, fibrotic processes typically extend into non-infarcted areas and alter cardiac structure.^31, 32^ However, post-infarction fibrotic processes were notably restricted in non-infarct regions of TKO hearts, coinciding with reduced expression of periostin and galectin-3 (Figure S3A,B).

#### Molecular Protection in Infarct Zones

Our RNA sequencing analysis of infarct territories revealed extensive preservation of mitochondrial gene expression in TKO hearts compared to wild-type controls (Figure 1G). Notably, all thirteen mitochondrial DNA-encoded genes showed significant preservation in TKO hearts, contrasting sharply with the widespread suppression observed in wild-type infarct tissue.^7, 33^ This pattern extended to nuclear genes affecting mitochondrial function, with maintained expression of genes involved in respiratory complexes and PGC-1α, a master regulator of mitochondrial biogenesis (Figure 1H).

The preservation of mitochondrial gene expression paralleled findings from pharmacological studies using the Parkin activator PR-364,^16^ suggesting a common mechanism involving enhanced mitochondrial quality control. Western blot analysis demonstrated remarkable preservation of cardiomyocyte-specific proteins within TKO infarct zones. Cardiac troponin I (cTnI), a highly specific cardiac marker, was markedly reduced in wild-type infarct zones but maintained at near-normal levels in TKO infarct areas. Similarly, SERCA2a protein, crucial for cardiac contractility through calcium handling regulation, showed dramatic preservation in TKO hearts (Figure 1I and Figure S4).

#### Enhanced Cardiomyocyte Survival

The preservation of both structural and functional proteins suggested enhanced cardiomyocyte survival within infarct territories. RNA sequencing quantification confirmed partially maintained expression of key cardiac genes including Myh6 (α-myosin heavy chain), Tnni3 (cardiac troponin I), Tnnc1 (cardiac troponin C), Tnnt2 (cardiac troponin T), Actn2 (α-actinin-2), Nkx2-5, and Atp2a2 (SERCA2a) in TKO infarct zones (Figure 1J). The preservation of these diverse cardiac markers indicated protection against multiple forms of cardiomyocyte death, including both apoptosis and necrosis. RNA-seq quantification also confirmed that ketolysis-related genes and fatty acid oxidation gene clusters were partially preserved in TKO infarct zones (Figure S5A,B).

### Parkin Upregulation as the Central Mediator of Cardioprotection

Our investigation of molecular mechanisms underlying cardiac protection revealed dramatic Parkin upregulation across multiple tissues in TKO mice, providing insight into the mechanistic basis of observed cardioprotection.

#### Tissue-Wide Parkin Induction

Quantitative RT-PCR analysis demonstrated significant increases in Parkin (Prkn) mRNA expression across multiple tissues. Both gastrocnemius muscle and heart tissue exhibited 6-fold increases (n=7-8, P<0.0001), while brain tissue showed 5-fold elevation (n=5, P<0.001) compared to wild-type controls (Figure 2A-C). The magnitude and consistency of these changes across diverse tissue types suggested fundamental alteration in Parkin gene regulation rather than tissue-specific adaptive responses.

**Figure 2.**
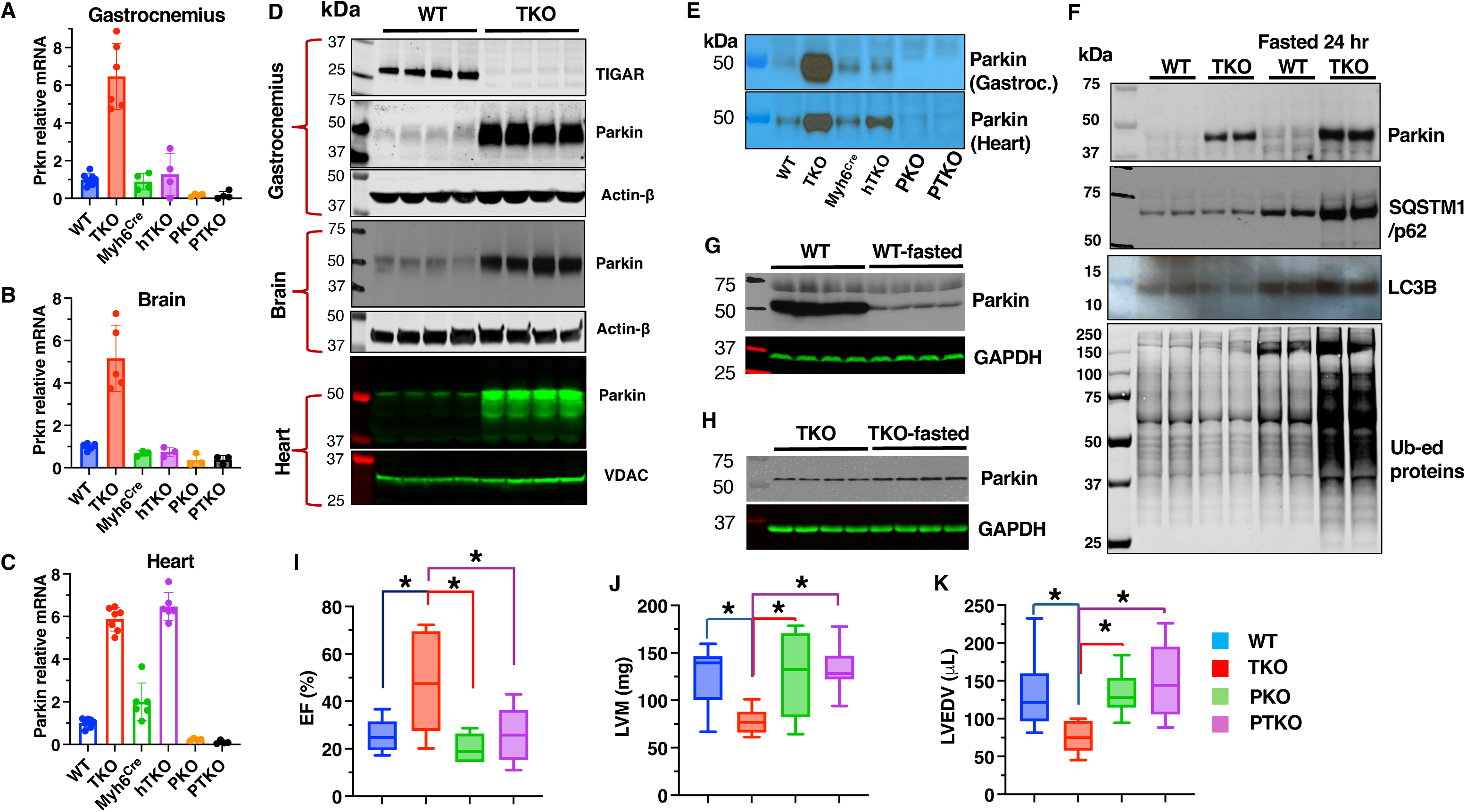
Parkin upregulation mediates cardiac protection in TKO and hTKO mice. A-C, Parkin (Prkn) mRNA levels in gastrocnemius muscle (A), brain (B), and heart (C) of WT, TKO, Myh6^Cre^, hTKO, Parkin knockout (PKO), and Parkin/TIGAR double knockout (PTKO) mice determined by qRT-PCR. Data normalized to WT expression. D, Western blot analysis of TIGAR and Parkin protein levels in gastrocnemius muscle, brain, and heart tissues from WT and TKO mice, with β-actin and VDAC as loading controls. E, Detection of Parkin protein in gastrocnemius muscle and heart tissues from WT, TKO, Myh6Cre, hTKO, PKO, and PTKO mice by immunoprecipitation (IP) followed by immunoblotting (IB). F, Western blot analysis of mitophagy-related proteins, including Parkin, SQSTM1/p62, LC3B, and ubiquitinated proteins in heart mitochondrial fractions from fed and 24-hour fasted WT and TKO mice. G-H, Immunoblot analysis of Parkin protein levels in cytosolic fractions from heart tissues of fed and 24-hour fasted WT (G) and TKO (H) mice, with GAPDH as loading control. I-K, Echocardiographic assessment of left ventricular ejection fraction (EF, I), mass (LVM, J), and end-diastolic volume (LVEDV) in 4-month-old male WT, TKO, PKO, and PTKO mice 4 weeks after MI. Data represent mean±SD; *P<0.05; n=5-10 per group.

Importantly, cardiomyocyte-specific TIGAR knockout mice (hTKO) also demonstrated significant Parkin upregulation in heart tissue (3-fold increase, n=6, P<0.0001) without corresponding changes in non-cardiac tissues such as gastrocnemius muscle or brain. This tissue-specific pattern in hTKO mice confirmed that TIGAR’s regulation of Parkin expression was cell-autonomous and did not require systemic TIGAR deficiency.

#### Protein-Level Confirmation

Western blot analyses confirmed these findings at the protein level, showing substantially elevated Parkin protein expression in TKO mice across all examined tissues (Figure 2D). Due to dramatic differences in Parkin protein levels between wild-type and TKO samples, immunoprecipitation studies were necessary to concentrate Parkin protein from wild-type samples for accurate quantitative comparisons (Figure 2E). These studies confirmed substantial upregulation in both TKO and hTKO hearts and skeletal muscle, but not in skeletal muscle from hTKO mice, consistent with mRNA expression patterns.

#### Enhanced Mitophagic Function

The functional significance of Parkin upregulation became evident during metabolic stress conditions. Under baseline fed conditions, Parkin phosphorylation at serine 65 (a marker of Parkin activation) was proportionally increased in TKO hearts relative to total Parkin levels, indicating functional competence of upregulated Parkin (Figure S6A).

During 24-hour fasting, TKO hearts exhibited enhanced mitophagic responses compared to wild-type controls, evidenced by increased mitochondrial protein ubiquitination, elevated recruitment of autophagy adaptor proteins (SQSTM1/p62), and enhanced LC3B lipidation (Figure 2F). These enhanced responses occurred without baseline depletion of mitochondrial proteins, indicating appropriate regulation without excessive mitochondrial loss (Figure S6B).

#### Preserved Mitophagic Capacity Under Stress

In the basal state, Parkin is primarily localized to the cytoplasm and translocates to the mitochondria when mitophagy becomes activated.^34^ In wild-type hearts, Parkin was depleted from the cytosolic fraction after 24 hours of fasting (Figure 2G), while TKO hearts maintained cytosolic Parkin reserves (Figure 2H). This resulted in greater mitochondrial-associated Parkin in both basal and fasted states in TKO hearts (Figure 2F). Despite increased Parkin protein levels, Parkin remains inactive until an appropriate cardiac mitochondrial stress is induced, then provides enhanced capacity for mitochondrial ubiquitination and mitophagy completion.

#### Genetic Confirmation of Parkin’s Essential Role

To definitively establish Parkin’s role in mediating cardioprotective effects, Parkin/TIGAR double knockout (PTKO) mice were generated. These animals showed complete loss of the protective phenotype previously observed in TKO mice, providing compelling genetic evidence for Parkin’s essential role.

Following myocardial infarction, PTKO mice exhibited severely impaired cardiac function, with post-infarction ejection fraction (25.77±11.61%, n=10) comparable to wild-type mice (25.32±7.25%, n=5) and significantly worse than TKO mice (46.94±20.47%, n=9, P=0.012) (Figure 2I). Similarly, parameters of left ventricular remodeling in PTKO mice closely resembled wild-type animals, with significantly increased left ventricular mass and end-diastolic volume compared to TKO mice but indistinguishable from wild-type controls (Figure 2J,K).

These findings provided definitive genetic proof that Parkin upregulation was both necessary and sufficient for cardioprotective effects conferred by TIGAR deficiency. The complete reversal of protection in PTKO mice eliminated the possibility that alternative pathways could compensate for Parkin’s absence in the setting of TIGAR deficiency.

### Comprehensive Protection Against Diet-Induced Metabolic Cardiomyopathy

Cardioprotective effects of TIGAR deficiency extended beyond acute ischemic injury to chronic metabolic stress conditions, providing evidence for broad-spectrum cardiac protection.

#### Resistance to Cardiac Hypertrophy

To model diet-induced metabolic cardiomyopathy, mice were subjected to a 6-month high-fat diet regimen containing 60% calories from fat. Wild-type mice developed significant cardiac hypertrophy under these conditions, as indicated by increased heart weight (182.7±17.2 mg, n=10) compared to chow-fed controls (137.9±6.4 mg, n=9). In striking contrast, TKO mice remained remarkably resistant to hypertrophic remodeling, showing only minimal cardiac mass increase (142±13.5 mg, n=10) compared to chow-fed TKO counterparts (126.8±6.4 mg, n=9) (Figure 3A).

**Figure 3.**
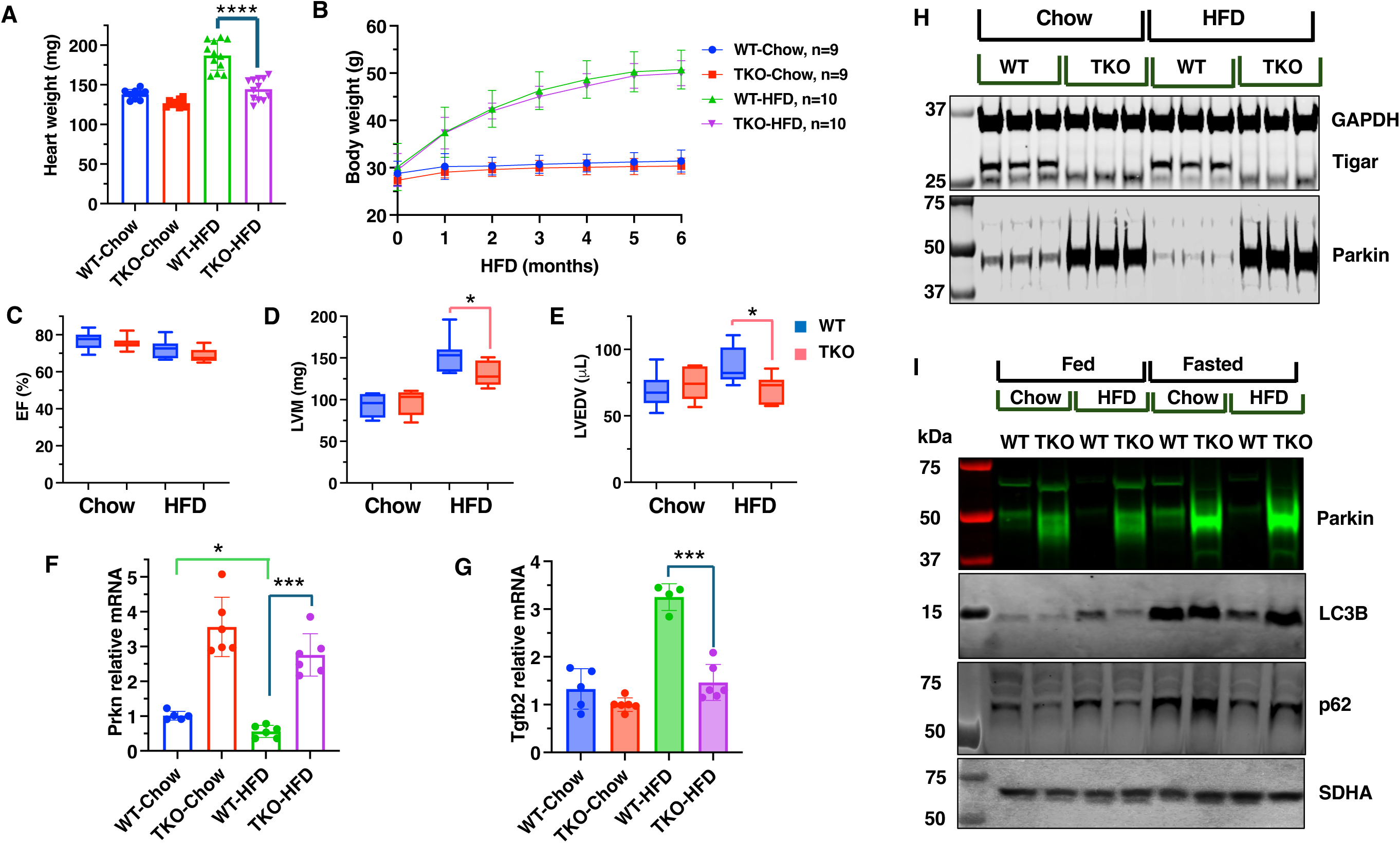
High-fat diet effects on cardiac function and mitophagy in WT and TKO mice. A, Heart weights of WT and TKO mice after six months of normal chow or high-fat diet (HFD). HFD increased cardiac mass in WT but not TKO mice. B, Body weight progression in WT and TKO mice during six months of normal chow or HFD (n=9-10 per group). C-E, Echocardiographic assessment of left ventricular ejection fraction (EF, C), mass (LVM, D), and end-diastolic volume (LVEDV, E) in WT and TKO mice fed normal chow or HFD. *<0.05. F-G, mRNA expression levels of Parkin (F) and TGF-β2 (G) in heart tissue from WT and TKO mice fed normal chow or HFD. Data normalized to WT-Chow expression. *P<0.05, ***P<0.001, n=4-6. H, Western blot analysis of TIGAR and Parkin protein levels in heart tissue cytosolic fraction from WT and TKO mice fed normal chow or HFD, with GAPDH as loading control. I, Western blot analysis of mitophagy-related proteins (Parkin, LC3B, and p62) in mitochondrial fractions of heart tissue from fed and 24-hour fasted WT and TKO mice maintained on normal chow or HFD. SDHA serves as mitochondrial fraction loading control. Data represent mean±SD.

Importantly, this protection occurred despite comparable body weight gains between genotypes (wild-type HFD: 50.3±5.0 g vs TKO HFD: 50.2±1.7 g, n=10), indicating that cardioprotective effects were independent of systemic metabolic load or differences in diet-induced obesity (Figure 3B).

#### Preserved Cardiac Geometry

Our echocardiographic analysis revealed that systolic function parameters remained preserved across all groups (Figure 3C), consistent with heart failure with preserved ejection fraction (HFpEF).^35^ Wild-type mice on high-fat diet exhibited pathological hypertrophic remodeling, including increased left ventricular mass (152.5±20.1 mg vs 131.0±14.2 mg in TKO, P<0.05) and elevated end-diastolic volume (87.5±13.67 μL vs 69.89±10.03 μL in TKO, P<0.05) (Figure 3D,E). TKO hearts maintained normal cardiac geometry, indicating substantial protection against pathological remodeling.

#### Molecular Mechanisms of Protection

Molecular analysis revealed distinct transcriptional and protein expression profiles corresponding to observed phenotypic differences. Wild-type mice subjected to high-fat diet showed significant downregulation of cardiac Parkin at both mRNA (Figure 3F) and protein levels (Figure 3H), consistent with previous reports of impaired mitochondrial quality control in metabolic cardiomyopathy.

This Parkin suppression was accompanied by upregulation of established pathological remodeling markers, including transforming growth factor beta-1 (Tgfb1), a key mediator of cardiac fibrosis and pathological remodeling (Figure 3G).^36^ TKO hearts maintained robust Parkin expression under high-fat diet conditions (Figure 3H) and showed no significant induction of these pathological markers, confirming protection against diet-induced cardiomyopathy at the molecular level (Figure 3D,E,F,G).

#### Preserved Mitophagic Responses

Our analysis of mitochondrial quality control mechanisms revealed critical functional differences between wild-type and TKO hearts under metabolic stress. Consistent with previous studies, wild-type hearts subjected to high-fat diet showed paradoxical increases in baseline autophagy markers (p62 and LC3B protein levels) despite substantial Parkin reduction. This elevation of autophagy markers suggests impaired mitophagic flux and accumulation, hallmarks of dysfunctional mitochondrial quality control (Figure 3I).

Critically, these hearts lost their adaptive mitophagic response to metabolic challenge, evidenced by failure to further upregulate p62 and LC3B protein levels in mitochondrial homogenates following 24-hour fasting compared to the fed state.^37, 38^ This impaired dynamic response indicated that Parkin deficiency had compromised the heart’s ability to respond appropriately to metabolic stress through enhanced mitochondrial quality control.^39–41^ In contrast, TKO hearts maintained a markedly different and healthier mitophagic profile under high-fat diet conditions. They exhibited low baseline levels of p62, suggesting efficient basal autophagy, while preserving robust Parkin expression even under metabolic stress. Most importantly, TKO hearts retained capacity for dynamic mitophagic responses to metabolic challenge, demonstrated by appropriate increases in both LC3B and p62 levels following 24-hour fasting, responses comparable to those observed under normal chow diet conditions (Figure 3I).

These findings demonstrated that sustained Parkin upregulation in TKO hearts provided comprehensive protection against both baseline mitochondrial dysfunction and loss of adaptive capacity that characterize diet-induced cardiomyopathy.

### Developmental Timing of TIGAR-Parkin Regulation

Having established that Parkin upregulation was essential for TIGAR deficiency-mediated cardioprotection, we investigated whether this relationship could be therapeutically exploited through adult interventions targeting TIGAR expression.

#### Adult TIGAR Manipulation Studies

To test whether suppressing TIGAR in adult hearts could elevate Parkin levels and recapitulate the cardioprotective phenotype, cardiac-specific adeno-associated virus (AAV9) vectors encoding wild-type TIGAR (TWT), a phosphatase-deficient TIGAR mutant (TMU), or TIGAR-specific short hairpin RNA (shTigar) were generated for selective expression in adult cardiomyocytes.

#### Re-expression Studies

Cardiac AAV9-mediated re-expression of wild-type TIGAR in adult TKO mice (TKO-TWT) successfully restored TIGAR mRNA levels to approximately 2-fold above wild-type levels, confirming effective viral transduction and transgene expression (Figure 4A). Similarly, the phosphatase-deficient mutant achieved 2.5-fold elevation above control levels. However, despite successful TIGAR protein restoration, neither wild-type nor phosphatase-deficient TIGAR re-expression affected the elevated Parkin mRNA or protein levels in TKO hearts (Figure 4B,E).

**Figure 4.**
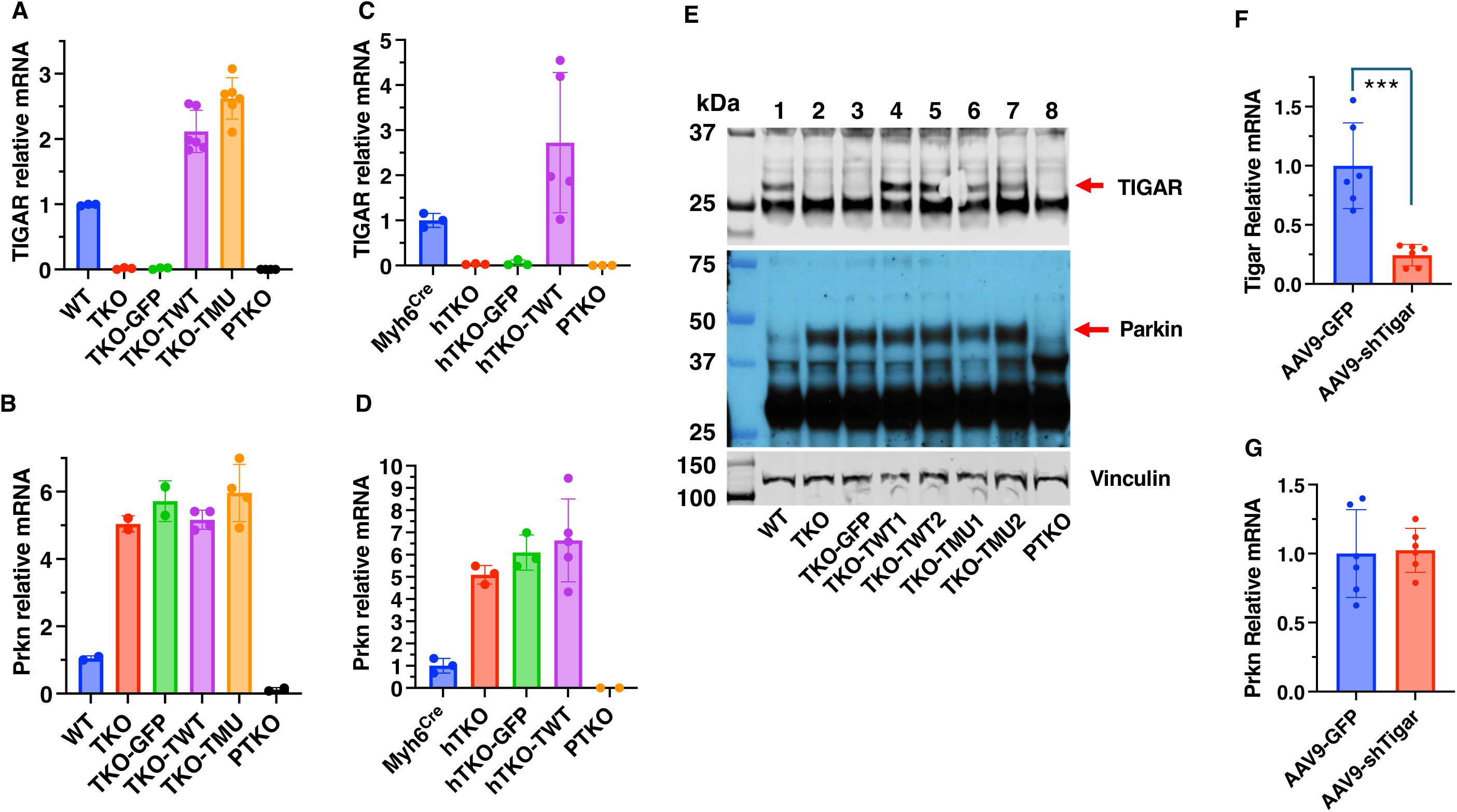
TIGAR expression in adult hearts does not directly affect Parkin expression. A-B, qPCR analysis of TIGAR (A) and Parkin (B) mRNA in heart samples from WT, TKO, TKO mice injected with AAV9-cTnT-GFP control virus (TKO-GFP), AAV9-cTnT-TIGAR wild-type virus (TKO-TWT), phosphatase-deficient TIGAR mutant virus (TKO-TMU), and PTKO mice. Data normalized to WT expression. C-D, qPCR analysis of TIGAR (C) and Parkin (D) mRNA in heart samples from Myh6^Cre^, hTKO, hTKO mice injected with AAV9-cTnT-GFP control virus (hTKO-GFP), AAV9-cTnT-TIGAR wild-type virus (hTKO-TWT), and Parkin-TIGAR double knockout (PTKO) mice. Data normalized to Myh6^Cre^ expression. E, Western blot analysis of TIGAR (30 kDa) and Parkin (52 kDa) protein expression in heart samples from various genotypes as indicated (lanes 1-8), with Vinculin (117 kDa) as loading control. Red arrows indicate TIGAR and Parkin bands. F-G, qPCR analysis of TIGAR (F) and Parkin (G) mRNA in heart samples from WT mice treated with AAV9-GFP control or AAV9-cTnT-TIGAR shRNA. Data represent mean±SD; ***P<0.001; n=5 per group.

This unexpected finding was reproduced in cardiomyocyte-specific knockout (hTKO) mice, where AAV9-mediated TIGAR overexpression achieved 2.5-fold increase in TIGAR mRNA compared to Myh6^Cre^ controls but failed to alter elevated Parkin expression levels characteristic of hTKO hearts (Figure 4C,D).

#### Knockdown Studies

Complementary experiments using AAV9-mediated cardiac-specific TIGAR silencing in adult wild-type mice successfully reduced TIGAR mRNA levels (76% reduction, P<0.001) but were unable to induce any increase in Parkin mRNA expression (Figure 4F,G). These findings were particularly striking given that genetic TIGAR deficiency from birth resulted in dramatic Parkin upregulation.

#### Implications for Developmental Programming

The failure of adult TIGAR manipulation to alter Parkin expression levels, despite successful modulation of TIGAR itself, suggested that TIGAR’s regulatory effect on Parkin occurs primarily during critical developmental windows rather than through ongoing adult regulation. Once established during development, the Parkin expression pattern appeared to become largely independent of continued TIGAR regulation.

This developmental timing aligned with known biology of cardiac maturation, where the transition from fetal to adult metabolism occurs postnatally and involves major reprogramming of mitochondrial function and quality control mechanisms. The findings suggested that TIGAR deficiency during this critical period establishes persistent alterations in Parkin gene regulation that are maintained throughout adult life.

### Tissue-Specific Regulation and Mechanistic Insights

To better understand the specificity and mechanisms of TIGAR-Parkin regulation, we conducted comprehensive analyses across multiple tissue types and cellular contexts.

#### Tissue-Specific Expression Patterns

TIGAR deficiency induced Parkin mRNA expression in heart, skeletal muscle, and brain (Figure 2A,B,C) but had no effect in testicular tissue, where wild-type mice already expressed relatively high baseline Parkin levels (Figure 5A,B). This tissue specificity was particularly evident when comparing isolated cardiomyocytes (CM) with testicular tissue from the same TKO animals.

**Figure 5.**
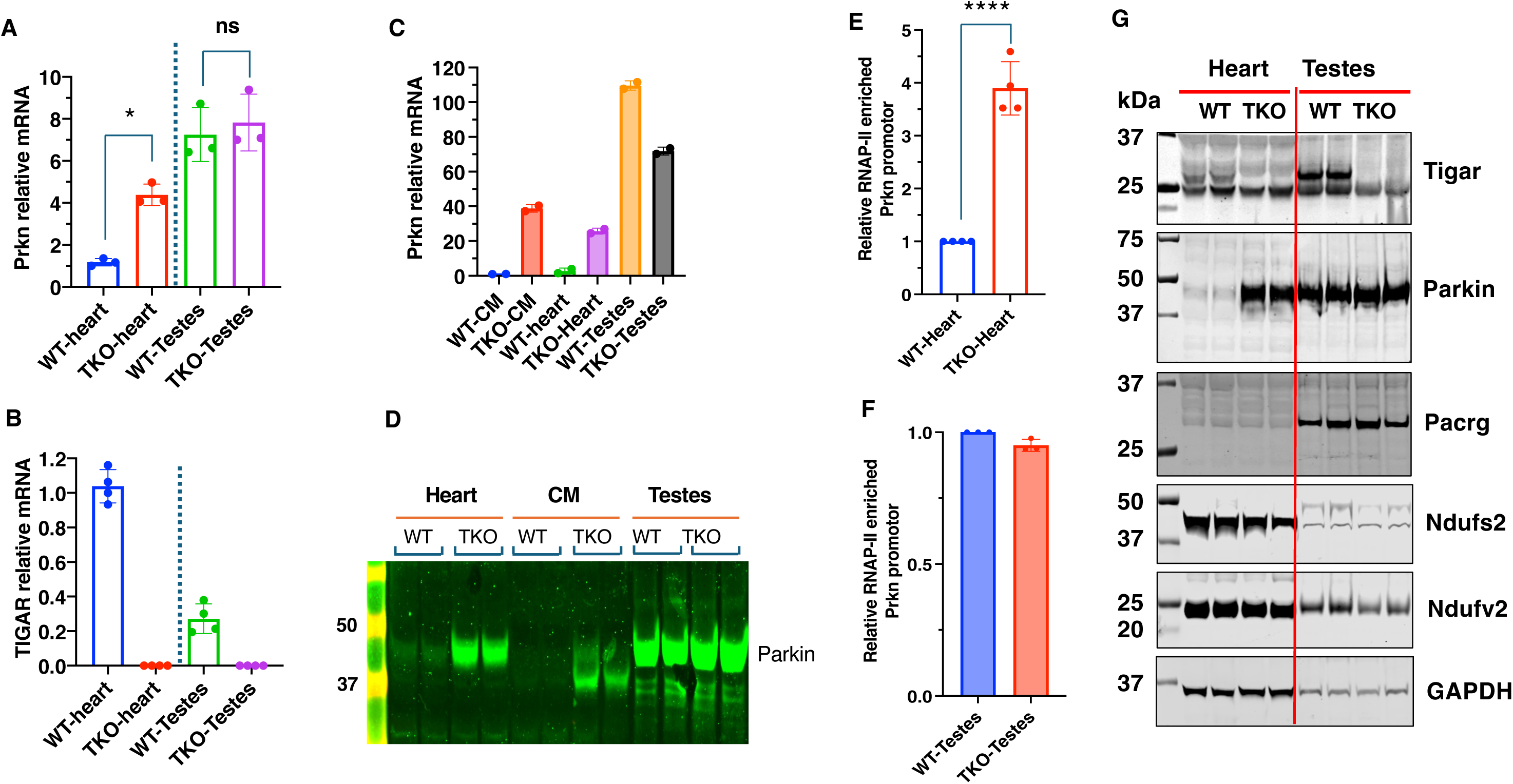
Tissue-specific regulation of TIGAR and Parkin expression in heart and testes. A-B, qPCR analysis of Parkin (A) and TIGAR (B) mRNA levels in heart and testes from WT and TKO mice. Data normalized to WT heart expression. Data represent mean±SD; ns, not significant; *P<0.05. C, qPCR analysis of Parkin mRNA levels in isolated fresh cardiomyocytes (CM), heart tissues, and testes tissues from WT and TKO mice. Data normalized to WT-CM expression. D, Western blot analysis of Parkin protein levels in heart tissues, isolated cardiomyocytes (CM), and testes tissues from WT and TKO mice. E-F, RNA polymerase II (RNAP-II) ChIP-qPCR analysis showing RNA polymerase II enrichment at the bidirectional Prkn/Pacrg promoter in WT and TKO heart tissue (E) and WT and TKO testes tissue (F). Data normalized to WT heart or testes and represent mean±SD; ***P<0.001; n=3 to 4 independent experiments. G, Western blot analysis of TIGAR, Parkin, Pacrg, Ndufs2, Ndufv2, and GAPDH protein levels in heart and testes from WT and TKO mice, demonstrating tissue-specific expression patterns.

Isolated TKO cardiomyocytes demonstrated a dramatic 39-fold increase in Parkin mRNA expression compared to wild-type cardiomyocytes, while testicular tissue maintained high baseline expression in both genotypes (Figure 5C,D). This cell-specific upregulation provided compelling evidence that enhanced Parkin expression resulted directly from TIGAR deficiency in a cell context-dependent manner.

#### Transcriptional Activation

ChIP-qPCR targeting RNA polymerase II at the Parkin promoter showed a 4-fold increase in occupancy in TKO cardiac tissue compared to wild-type controls (P<0.0001), indicating enhanced transcriptional activation (Figure 5E). Testicular tissue showed no difference between genotypes (Figure 5F). Parkin shares a 203 bp promoter bidirectional promoter that also controls the expression of the Pacrg gene.^42^ TKO hearts displayed robust Parkin protein induction but not Pacrg protein, suggesting TIGAR’s influence involved regulatory elements beyond the shared promoter region (Figure 5G). As controls the mitochondria respirator chain subunits, Ndufs2 and Ndufv2 respirator chain subunits and cytosolic glycolytic protein GAPDH were unaffected in either the heart or testes of TKO mice.

### Epigenetic Mechanisms of Parkin Gene Regulation

The developmental timing of TIGAR’s effects on Parkin expression, combined with persistence of elevated Parkin levels despite adult TIGAR restoration, strongly suggest developmental epigenetic mechanisms underlying this regulatory relationship. Since epigenetic histone modifications tend to be relatively reversible whereas DNA cytosine methylation (5mC) is relatively stable, we speculated that the methylation of the Parkin gene may underlie the presistent TIGAR-dependent changes in Parkin transcription.

#### Whole-Genome Bisulfite Sequencing

We performed comprehensive WGBS analysis of cardiac and testicular tissue from wild-type and TKO mice to assess DNA methylation patterns and identify differentially methylated regions (DMRs) associated with TIGAR deficiency. RNA sequencing tracks demonstrated increased Parkin transcript levels across all 12 exons in TKO hearts (Figure 6A,B). WGBS analysis identified multiple DMRs throughout the Parkin gene body with consistently lower methylation levels in TKO hearts.

**Figure 6.**
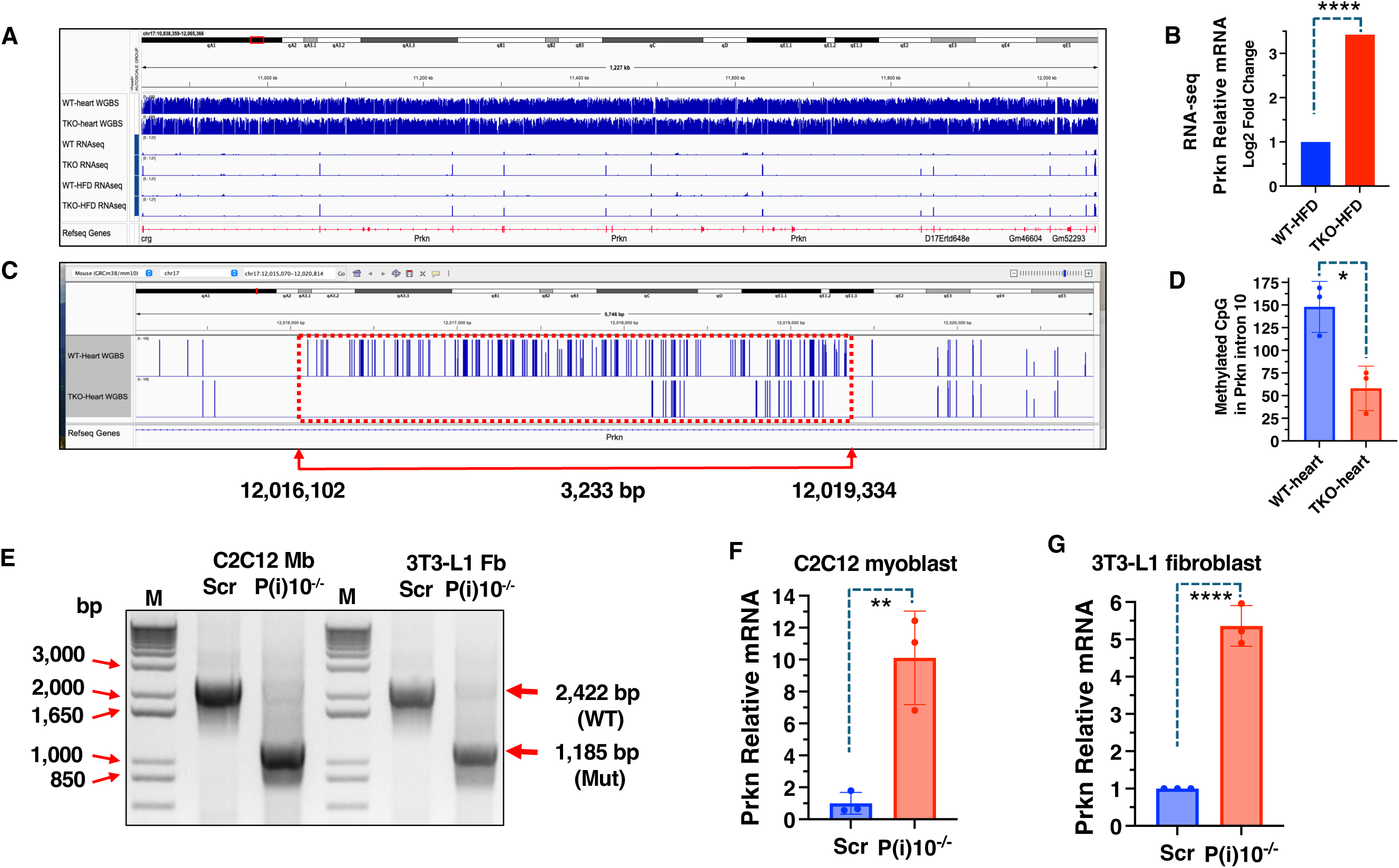
Prkn gene body methylation patterns and Parkin expression regulation. A, Integrative Genomics Viewer (IGV) visualization of the Prkn gene with whole-genome bisulfite sequencing (WGBS) and RNA-seq data from WT and TKO mice hearts under normal chow and HFD conditions, with corresponding RNA-seq analysis showing significantly increased Parkin (Prkn) mRNA expression in TKO-HFD vs WT-HFD hearts (B). C, Enlarged view of the differentially methylated region (DMR) within Prkn intron 10 (red dashed box, coordinates chr17:12,016,102-12,019,334, 3,233 bp). D, Quantification of methylated CpG sites in the DMR in Prkn intron 10 in WT (148±28) and TKO (58±24) heart tissues. *P=0.0139, n=3. E, CRISPR/Cas9-mediated deletion of a 14,177 bp fragment (chr17:12006677 to chr17:12020853) in Prkn intron 10 in C2C12 myoblasts (Mb) and 3T3-L1 fibroblasts (Fb). Agarose gel electrophoresis shows the predicted 1,185 bp PCR amplicon in P(i)10^-/-^ cells (gRNA/Cas9 vector transfected) vs the 2,422 bp band in scramble vector (Scr) transfected control cells. F-G, RT-qPCR analysis of Parkin mRNA levels in Prkn intron 10 DMR-deleted (P(i)10^-/-)^ and scramble control (Scr) C2C12 myoblasts (F) and 3T3-L1 fibroblasts (G), showing 10-fold and 5-fold increases, respectively, in Parkin expression following DMR deletion. Data represent mean±SD; **P<0.01; ***P<0.001; n=3 independent experiments.

#### Identification of Regulatory DMR

Among identified DMRs, a 3.2 kb region within Parkin intron 10 showed particularly significant hypomethylation in TKO hearts (Figure 6C). Quantitative analysis revealed that average methylation levels in this region were substantially reduced in TKO compared to wild-type hearts (58±24 vs 148±28 methylated CpG sites, n=3, P=0.0139) (Figure 6D).

This intronic DMR was particularly intriguing given its location within the gene body rather than the promoter region. While the Parkin promoter region contains a CpG island, no significant methylation changes were detected in this region, suggesting that TIGAR’s regulatory effects were mediated through alternative epigenetic mechanisms.

#### Functional Validation Through Gene Editing

We employed CRISPR/Cas9-mediated deletion of a 14.2 kb region (chr17:12006677-12020853) encompassing the 3.2 kb DMR using dual guide RNAs (Figure 6E). PCR analysis confirmed successful deletion in both C2C12 myoblasts and 3T3-L1 fibroblasts, with mutant samples showing the expected 1,185 bp amplicon compared to the 2,422 bp product in control cells. Quantitative RT-PCR analysis demonstrated that this deletion dramatically enhanced Parkin mRNA expression in both cell types, with an approximately 10-fold increase in C2C12 myoblasts (Figure 6F) and 6-fold increase in 3T3-L1 fibroblasts (Figure 6G) compared to scramble controls. These findings provided direct functional evidence that the identified DMR normally functions as a negative regulatory element.

## DISCUSSION

This study reveals a novel mechanism whereby TIGAR deficiency establishes persistent cardioprotection through developmental epigenetic programming of Parkin expression. This regulatory relationship confers broad protection against both acute ischemic injury and chronic metabolic cardiomyopathy through enhanced mitochondrial quality control mechanisms established during cardiac development and maintained throughout life.

### A Novel TIGAR-Parkin Regulatory Relationship

Our discovery of TIGAR-Parkin regulatory interaction represents a significant advancement in understanding cardiac metabolic programming and mitochondrial quality control. Previous studies demonstrated cardioprotection in TIGAR knockout mice against ischemia and pressure overload heart failure.^26–28^ However, the molecular mechanisms underlying this protection remained unclear. These findings reveal that TIGAR’s protective effects depend critically on developmental timing and connection to Parkin expression.

The dramatic increase in Parkin expression observed across multiple tissues in TIGAR-deficient mice (6-fold in heart and skeletal muscle, 5-fold in brain) represents one of the most striking examples of metabolic gene regulation encountered in cardiac research. The consistency of this effect across diverse tissues, combined with heart-specific knockout models, indicates this is a fundamental regulatory relationship rather than tissue-specific adaptive responses.

Genetic rescue experiments using Parkin/TIGAR double knockout mice, which eliminated cardioprotection, provided definitive genetic evidence for Parkin’s essential role. This approach represents convincing method to prove causality in mouse models and rules out alternative explanations for observed phenomena.

### Developmental Programming of Cardiac Resilience

One of the most striking aspects of our findings relates to how developmental programming shapes cardiovascular health throughout life. The fact that adult TIGAR manipulation could not alter Parkin expression levels, despite successfully changing TIGAR itself, demonstrates that this regulatory relationship becomes established during critical developmental windows and remains fixed in adult tissues.

This developmental timing aligns with the well-characterized postnatal shift in cardiac metabolism from glycolysis-dependent fetal metabolism to fatty acid oxidation-dependent adult metabolism.^19^ During this transition, cardiomyocytes undergo massive mitochondrial expansion, maturation of respiratory complexes, and establishment of quality control mechanisms that serve the heart throughout adult life.^5, 18, 19^

Our findings suggest that metabolic conditions during critical developmental periods can permanently alter expression of key mitochondrial quality control genes through epigenetic mechanisms, with profound implications for understanding how early environmental influences affect lifelong cardiovascular health. These discoveries highlight the critical importance of optimal maternal and childhood nutrition in programming cardiac resilience, suggesting that nutritional interventions during early development could establish lifelong protection against cardiovascular disease through enhanced mitochondrial quality control pathways.

### Epigenetic Mechanisms and Therapeutic Implications

Our identification of a specific 3.2 kb differentially methylated region within Parkin intron 10 that controls gene expression provides important mechanistic insights and potential therapeutic targets. Unlike promoter-proximal regulatory elements that control transcriptional initiation,^43^ this intronic region likely influences transcriptional elongation efficiency through the large Parkin gene.

Functional validation of this regulatory element through CRISPR-mediated deletion, which enhanced Parkin expression 6-10 fold in multiple cell types, demonstrates its potential as a therapeutic target. Future approaches might involve targeted demethylation of this region using emerging epigenome editing technologies, potentially allowing manipulation of Parkin expression levels in adults.

The tissue-specific nature of TIGAR’s effects on Parkin expression is noteworthy, testicular tissue remained completely unresponsive despite high baseline Parkin levels. This suggests tissue-specific epigenetic regulatory mechanisms that require further investigation. Understanding these could enable tissue-targeted therapeutic interventions that enhance cardiac Parkin expression without affecting other organ systems.

### Clinical Relevance and Translational Potential

These findings have several important clinical implications extending beyond basic mechanistic insights:

#### Ischemic Heart Disease

The remarkable preservation of cardiac function following myocardial infarction in TIGAR-deficient mice (maintaining 60% vs 36% of baseline function) represents a degree of cardioprotection that would be clinically transformative if achievable in human patients. Enhanced survival of cardiomyocytes within damaged zones, demonstrated by preservation of cardiac troponin I, SERCA2a, and other essential proteins, suggests that enhanced Parkin expression could extend the window for reperfusion therapy and improve outcomes even when treatment is delayed.^44, 45^

#### Diabetic Heart Disease

The complete resistance to diet-induced cardiac dysfunction observed in TKO mice, despite equivalent weight gain, directly addresses one of the major challenges in managing diabetic cardiovascular disease. Current therapies often fail to prevent cardiac complications even with optimal glycemic control, highlighting the need for approaches targeting underlying mitochondrial dysfunction.^8, 46^ The preservation of normal cardiac structure and mitophagic responses in TKO hearts suggests that enhancing Parkin expression could prevent diabetic heart disease development.^39–41^

#### Heart Failure with Preserved Ejection Fraction

The dysfunction observed in wild-type mice on high-fat diet closely mirrors the clinical syndrome of heart failure with preserved ejection fraction (HFpEF), which represents approximately half of all heart failure cases and currently lacks effective therapies.^47^ The protection against this condition in TKO mice suggests that targeting mitochondrial quality control could provide new therapeutic approaches for this challenging clinical problem.

### Mechanistic Insights into Mitochondrial Quality Control

Our study provides important insights into mitochondrial quality control regulation and function in the heart. The enhanced mitophagic responses observed in TKO hearts during metabolic stress, combined with preservation of cytosolic Parkin reserves even under prolonged stress conditions, demonstrate that sustained high-level Parkin expression enhances cardiac adaptive capacity without compromising baseline mitochondrial function.

The preservation of normal mitochondrial protein levels under baseline conditions, despite enhanced mitophagic machinery, indicates that upregulated Parkin does not result in excessive mitochondrial degradation. This suggests that mitophagy is appropriately regulated and responds primarily to damaged or dysfunctional organelles rather than randomly targeting healthy mitochondria.

The loss of adaptive mitophagic responses in wild-type hearts subjected to high-fat diet, demonstrated by failure to upregulate autophagy markers during fasting despite metabolic stress, highlights how mitochondrial quality control becomes progressively impaired in metabolic disease.^48^ The preservation of these responses in TKO hearts demonstrates that enhanced Parkin expression can prevent this deterioration and maintain cardiac adaptive capacity under adverse conditions.^49^

### Comparison with Pharmacological Approaches

Our findings complement and extend recent studies using pharmacological Parkin activators such as PR-364, which demonstrated cardioprotection following myocardial infarction.^16^ The preservation of mitochondrial gene expression in TKO infarct zones closely matched observations with PR-364 treatment, suggesting common mechanisms involving enhanced mitochondrial quality control.

However, the genetic approach provides several advantages over pharmacological intervention. First, it establishes that beneficial effects result specifically from enhanced Parkin expression rather than off-target drug effects. Second, it demonstrates that sustained elevation of Parkin expression is well-tolerated and beneficial throughout the lifespan. Third, it reveals the importance of developmental timing in establishing effective cardioprotection.

### Limitations and Future Directions

These investigations were conducted in mouse models, and translation to human physiology will require validation in human tissues and clinical studies. Additionally, optimal methods for therapeutic intervention targeting this pathway remain to be determined, and potential long-term consequences of sustained Parkin elevation require additional investigation. Future studies should focus on developing approaches to target our identified epigenetic regulatory elements in adult animals, investigating precise metabolic pathways linking TIGAR deficiency to DNA methylation changes, and assessing relevance of these findings in human populations.

### Broader Implications for Cardiovascular Medicine

The concept of developmental programming revealed by this study has broader implications for cardiovascular medicine beyond the specific TIGAR-Parkin connection. It suggests that cardiac responses to adult stressors are fundamentally shaped by developmental experiences and that interventions during critical developmental windows could have lifelong beneficial effects.

This concept is particularly relevant given growing recognition that cardiovascular disease often begins early in life, even when clinical manifestations appear decades later.^50, 51^ These findings suggest that identifying and targeting developmental programming mechanisms could provide new approaches for preventing cardiovascular disease before it begins. Furthermore, the identification of specific epigenetic regulatory elements provides a framework for understanding how environmental factors during development can permanently alter disease susceptibility. This knowledge could inform recommendations for maternal and early childhood health interventions aimed at optimizing cardiovascular development.

### Conclusions

Our study identifies a novel TIGAR-Parkin regulatory pathway operating through epigenetic mechanisms during cardiac development to establish lifelong cardioprotection. TIGAR deficiency enhances Parkin expression through reduced methylation of a specific regulatory region in the Parkin gene, resulting in improved mitochondrial quality control and enhanced cardiac resilience against both myocardial infarction and metabolic stress. These findings provide new therapeutic targets for cardiovascular disease and highlight how developmental programming mechanisms determine lifelong cardiovascular health, opening new avenues for both prevention and treatment of heart disease.

## Nonstandard Abbreviation and Acrronyms

TIGAR: TP53-induced glycolysis and apoptosis regulator
CRISPR: Clustered Regularly Interspaced Short Palindromic Repeats
TKO: Tigar whole-body knockout
hTKO: Cardiomyocyte-specific knockout
DAMPs: mitochondrial damage-associated molecular patterns
PINK1: PTEN-induced kinase 1
TFAM: Transcription Factor A, Mitochondrial
PGC-1α: Peroxisome proliferator-activated receptor gamma coactivator 1-alpha
PKO: Parkin knockout
PTKO: Parkin/TIGAR double knockout
WGBS: Whole-genome bisulfite sequencing
ChIP: Chromatin immunoprecipitation
Cas9: CRISPR-associated protein 9
cTnI: Cardiac troponin I
PR-364: a small molecule that activates the protein Parkin
Myh6: Cardiac α-myosin heavy chain
Tnni3: troponin I3, cardiac type
Tnnc1: Troponin C1, slow skeletal and cardiac type
Tnnt2: Cardiac troponin T
Actn2: α-actinin-2
Nkx2-5: NK2 homeobox 5
Atp2a2: Sarco(endo)plasmic reticulum calcium-ATPase 2 (SERCA2)
RNA-seq: RNA sequencing
Prkn: Parkin gene SQSTM1/p62, Sequestosome 1
LC3B: Microtubule-associated protein 1 light chain 3 beta
HFpEF: heart failure with preserved ejection fraction
Tgfb1: Transforming growth factor beta-1
AAV9: Adeno-associated virus serotype 9
TWT: wild-type TIGAR
TMU: phosphatase-deficient TIGAR mutant
Myh6^Cre^: Transgenic mouse line where the Cre recombinase gene is expressed under the control of the Myh6 promoter
Tigar^fl/fl^: Transgenic mouse line where the Tigar gene is flanked by loxP sites
CM: Isolated cardiomyocyte
Pacrg: Parkin Co-Regulated Gene
Ndufs2: NADH dehydrogenase [ubiquinone] iron-sulfur protein 2, mitochondrial
Ndufv2: NADH dehydrogenase [ubiquinone] flavoprotein 2, mitochondrial.
GAPDH: Glyceraldehyde-3-phosphate dehydrogenase
DMRs: Differentially methylated regions
CpG island: cytosine-phosphate-guanine islands
LAD: Left anterior descending (LAD) artery ligation
EF: Left ventricular ejection fraction
LVM: Left ventricular mass
LVEDV: Left ventricular end-diastolic volume
HFD: High-fat diet
RNAP-II: RNA polymerase II

## ACKNOWLEDGMENTS

The authors thank members of the Pessin and Santulli laboratories for helpful discussions and technical assistance. Special thanks to Ms. Shiori Okada for technical support and Dr. Victor Schuster (Albert Einstein College of Medicine) for critical scientific discussion and project guidance.

## SOURCES OF FUNDING

This study was supported by grants DK033823, DK020541, and HL146691 from the National Institutes of Health.

## DISCLOSURES

The authors declare no competing financial interests.

## SUPPLEMENTAL MATERIAL

### Detailed Methods and Materials Animal Studies and Tissue Collection

All procedures were conducted under protocols approved by the Einstein Institutional Animal Care and Use Committee. Mice were housed in a facility with a 12-hour light/dark cycle and maintained on normal chow diet (5053, LabDiet) containing 62.3% carbohydrates, 24.5% protein, and 13.1% fat by caloric content. TKO and Tigarfl/fl mice were produced as described in our previous paper ^1^. The Tigar fl/fl mice were generated from C57BL/6N-Tigartm1a(EUCOMM)Wtsi/Wtsi mice obtained from the Wellcome Trust Sanger Institute (Hinxton Cambridge, UK) ^1^. Heart cardiomyocyte-specific Tigar knockout (hTKO) mice were produced by crossing Tigar fl/fl mice with cardiac-specific alpha myosin-heavy chain Cre (Myh6Cre) mice (The Jackson Laboratory, stock #011038). Genotyping was performed using primers 9543, 9544, oIMR8744, and oIMR8745 for Myh6Cre, and Tigarfl/fl-specific primers ^1^. for hTKO mice. Parkin knockout (PKO) mice were purchased from The Jackson Laboratory (Stock # 006582). Parkin and Tigar double knockout (PTKO) mice were produced by crossing PKO with TKO. At 16 weeks of age, mice underwent left anterior descending coronary artery ligation and echocardiographic imaging.

### Mouse heart tissue collection and processing for genomic DNA, total RNA, and protein isolation

Adult mice were briefly anesthetized in an induction chamber with 3-4% vaporized isoflurane (VETEQUIP, Pleasanton, CA) until loss of righting reflex, followed by cervical dislocation following institutional guidelines and approved protocols. Hearts were rapidly excised via thoracotomy and immediately placed in ice-cold phosphate-buffered saline (PBS, pH 7.4) to remove residual blood. The atria were carefully removed and the ventricles quickly blotted dry with lint-free tissue paper.

The ventricles were immediately snap-frozen using a cryogenic tissue clamp pre-chilled in liquid nitrogen. Frozen cardiac tissue was then transferred to a mortar immersed in liquid nitrogen and homogenized to a fine powder with a pre-chilled pestle, maintaining cryogenic conditions throughout the pulverization process to prevent tissue thawing. The tissue powder was collected into pre-labeled, sterile cryovials and stored at −80°C until further analysis.

### Total RNA extraction and quantitative RT-PCR

Total RNA from cells and tissues was extracted using the Direct-zol RNA MiniPrep Plus kit (Zymo Research) following the manufacturer’s instructions. First-strand cDNA was synthesized using the SuperScript IV VILO cDNA synthesis kit (Thermo Fisher Scientific). Quantitative TaqMan RT-PCR was performed on a QuantStudio 6 Flex platform (Applied Biosystems) to measure target mRNA expression. Relative gene expression was calculated using the ΔΔCt method and normalized to the internal control gene RNA18S1 or Rplp0. Quantitative RT-PCR results were analyzed using QuantStudio Design & Analysis Software (Applied Biosystems). The TaqMan primer-probe assays used in this study are listed below:

| Gene name | Gene symbol | TaqMan assay ID |
| --- | --- | --- |
| TP53-inducible Glycolysis and Apoptosis Regulator | Tigar | Mm00621530_m1 |
| Transforming growth factor β-2 | Tgfb2 | Mm00436955_m1 |
| Cardiac troponin T | Tnnt2 | Mm01290256_m1 |
| Parkin | Park2 | Mm01323528_m1 |
| Eukaryotic 18S rRNA Endogenous Control | RNA18S1 | 4310893E |
| Ribosomal protein, large, P0 | Rplp0 | Mm00725448_s1 |

For detection of mutant TIGAR (H11A/E102A/H198A) expression, custom primer-probe sets were used. The Tigar primer-probe set spanning exons 3-4 (PrimeTime Std qPCR Assay, Mm.PT.56a.16927616, 9630033F20Rik, Integrated DNA Technologies) and the internal control Rplp0 primer-probe set spanning exons 5-6 (Mm.PT.58.43894205) were used with PrimeTime Gene Expression Master Mix (Cat# 1055772, IDT) following the manufacturer’s protocol.

### Immunoblotting

Samples were prepared from cultured cells (washed with cold PBS) or tissues by homogenization with radioimmune precipitation assay (RIPA) lysis buffer (sc-24948, Santa Cruz Biotechnology) containing Halt protease and phosphatase inhibitor mixture (Cat# 78442, Thermo Fisher Scientific), 20 μM MG132, and 20 μM ALLN (EMD Millipore, Darmstadt, Germany) using Ceria stabilized zirconium oxide beads (MidSci, Valley Park, MO). Homogenates were centrifuged for 15 min at 21,000 × g at 4°C and supernatants collected for protein quantification using the BCA method. Protein samples were separated by either self-cast SDS-PAGE or SurePAGE (Bis-Tris) precast gel (GenScript, Piscataway, NJ) and transferred to nitrocellulose membrane using iBlot 2 Blotting System (Thermo Fisher Scientific). The immunoblot membrane was blocked with Pierce Protein-Free T20 (TBS) blocking buffer (product no. 37571, Thermo Fisher Scientific) or 6% milk in TBS with Tween 20 (TBST), then incubated with the primary antibody in blocking buffer or 1% milk TBST. Blots were washed in TBST and incubated with either IRDye 800CW secondary antibody (LI-COR, Lincoln, NE) or horseradish peroxidase-conjugated secondary antibody in blocking buffer. The membrane was washed with TBST and visualized using either the Odyssey CLx Imaging System (LI-COR) or enhanced chemiluminescence (ECL) (Thermo Fisher Scientific Pierce) method. Commercial primary antibodies were purchased from the following sources: Santa Cruz Biotechnology: TIGAR (sc-677290), SERCA2 (sc-376235), Parkin (sc-32282), Tom70 (sc-390545), NDuFS2 (sc-390596), NDuFV2 (sc-271620), vinculin (sc-73614), SDHA (sc-166909 HRP), SQSTM1/p62 (sc-48402), MAP LC3b (sc-376404), and β-actin (sc-47778). Cell Signaling Technology: Parkin (2132S), ParkinSer65 (36866S), troponin I (13083S), VDAC (4866S), Ubiquitin (43124S), and α-actinin 1 (6487S). ThermoFisher Scientific: Parkin (#702785) and PACRG (PA5-110069). EMD Millipore: Complex I 75-kDa (ABN302). Cosmo Bio USA: GAPDH (MBL-M171-3).

### Chromatin Immunoprecipitation and Quantitative PCR Analysis

Left ventricular tissue (approximately 40 mg) was collected from WT and TKO mouse hearts, washed with ice-cold PBS to remove blood, and immediately minced on an ice-embedded Petri dish using a pre-chilled razor blade. The minced tissue was cross-linked with 1% formaldehyde (ThermoFisher Scientific, #28908) and processed for chromatin isolation using the ChromaFlash™ Chromatin Extraction Kit (EpigenTek, P-2001). Chromatin samples (300 μl per 1.5 ml tube) were sonicated using a Diagenode Bioruptor for 15 cycles (30 seconds on/30 seconds off) at high power setting to generate DNA fragments ranging from 150-1000 base pairs. Chromatin immunoprecipitation was performed using the ChromaFlash™ One-Step ChIP Kit (EpigenTek, P-2025) following manufacturer’s instructions. DNA-protein complexes were immunoprecipitated using anti-RNA polymerase II monoclonal antibody ([CTD4H8], EpigenTek, A-2032-100), with non-immune IgG (EpigenTek, P-2025) serving as a negative control. The enriched DNA fragments and input DNA (10% of sample chromatin) were purified, released, and eluted. DNA concentration was determined using the Qubit dsDNA High Sensitivity Quantification Assay (ThermoFisher Scientific). Quantitative PCR analysis was performed using primers targeting the Prkn/Pacrg bi-directional promoter (Forward: GTCAACATTAGGAGACGCTAGTC; Reverse: GCAACTGTCTTCGCTGGTA) to generate a 76 bp amplicon, with mouse RPL30 intron 2 primers (Cell Signaling Technology, #7015P) serving as an internal control. PCR reactions were conducted using PowerUp™ SYBR™ Green Master Mix (ThermoFisher Scientific, A25741) on a QuantStudio 6 Flex platform. ChIP-qPCR data were analyzed using the 2–ΔΔ CT method.

### Myocardial infarction model and cardiac function assessment

Male mice (16-week old) underwent permanent ligation of the left anterior descending (LAD) coronary artery to induce MI ^2, 3^. Following baseline cardiac assessment, a left thoracotomy (1.5 cm) was performed through the fourth intercostal space under appropriate anesthesia. After retracting the lungs and opening the pericardium to expose the heart, the LAD coronary artery was identified and ligated at its proximal region using 8-0 silk suture. The ligation site was positioned between the pulmonary outflow tract and left atrial edge, approximately 2 mm distal to the left auricle tip. Successful arterial occlusion was confirmed by visible pallor of the anterior left ventricular (LV) wall. Post-operative care included maintaining body temperature using a heating pad until full recovery. Sham-operated controls underwent identical surgical procedures without arterial ligation.

Cardiac function was evaluated via transthoracic two-dimensional echocardiography using a 12-MHz probe (VeVo; Visualsonics) at baseline and weeks 1, 2, and 4 post-surgeries. LV parameters were measured through M-mode imaging in the parasternal short-axis view, including end-diastolic and end-systolic dimensions, septal and posterior wall thicknesses, to calculate fractional shortening and ejection fraction.

### Adult Mouse Cardiomyocyte Isolation

Cardiomyocytes were isolated as previously described ^2, 4^. Adult mice were injected intraperitoneally with 2,000 IU/kg heparin 5 min before heart isolation. Hearts were isolated and perfused using a Langendorff apparatus. Hearts were initially retroperfused for 20 minutes with calcium-free cardiac cell isolation buffer (containing in mM: NaCl: 113, KCl: 4.7, NaH₂PO₄: 0.6, KH₂PO₄: 0.6, HEPES: 10, Glucose: 7, Taurine: 15, 2,3-butanedione monoxime: 10, MgSO4: 1.2; pH 7.4) supplemented with 5 mM EDTA at 37°C. This was followed by enzymatic digestion through retroperfusion with the same buffer containing 300 U/mL collagenase type 4 (CLS-4, Worthington Biochemical Corp, Worthington, NJ) and 1.2 U/mL Protease from Streptomyces griseus type XIV (P5147, Sigma) for 30 minutes.

After perfusion, the left ventricle was carefully dissected and minced into approximately 1 mm³ pieces in cardiac cell isolation buffer containing collagenase II (300 U/mL) and protease XIV, then gently dissociated using forceps. The enzymatic digestion was terminated after 10 minutes by adding cardiac cell isolation buffer supplemented with 5% sterile, exosome-free FBS. The cell suspension was filtered through a 100-μm pore-size nylon mesh filter (22363549, Fisherbrand) to remove undigested tissue.

Isolated cells were allowed to settle by gravity in 15 mL tubes for approximately 20 minutes. During this sedimentation period, calcium was gradually reintroduced in four sequential steps (at 4-minute intervals) to reach final concentrations of 0.06, 0.24, 0.6, and 1.2 mM, respectively. Rod-shaped cardiomyocytes settled to form a pellet at the bottom of the tube and were collected for subsequent experiments. The quality and morphology of isolated cardiomyocytes were confirmed by microscopic examination.

Cardiomyocyte total RNA was extracted using Quick-RNA Miniprep (Zymo Research Inc., R1055) for Prkn mRNA quantification by qPCR. Total protein of cardiomyocytes was isolated using the Total Protein Extraction Kit for Animal Cultured Cells and Tissues (Invent Biotechnologies Inc., SD-001/SN-002), followed by Western blot analysis of Parkin protein expression as described previously.

### Tissue collection for RNA, protein, and immunofluorescence analyses post-LAD-MI

For RNA and protein analyses, the infarct, border, and remote zones were isolated, immediately snap-frozen in liquid nitrogen, pulverized in liquid nitrogen, and stored at - 80°C for subsequent RNA and protein extraction. For immunofluorescence studies, fresh cardiac tissues were embedded in Tissue-Tek O.C.T. compound and sectioned to 5 μm thickness. Immunostaining was performed using anti-periostin (PA5-34641, Invitrogen) and anti-α-actinin (sc-17829 AF488, Santa Cruz) antibodies. Stained sections were visualized and images captured using an Echo Revolve Microscope (San Diego, CA).

### Subcellular fractionation of mouse heart tissue for isolation of cytosolic and mitochondrial components

Mouse hearts were dissected from isoflurane-anesthetized animals and washed with ice-cold mitochondria assay solution (MAS; 70 mM Sucrose, 220 mM D-Mannitol, 5 mM KH2PO4, 5 mM MgCl2, 1 mM EGTA, 2 mM HEPES, 0.025% fatty acid free BSA, pH 7.4 adjusted with KOH) to remove blood. The heart was placed on an ice-embedded petri dish and finely minced using a pre-chilled razor blade. The minced heart tissue was homogenized in cold MAS buffer with Halt protease and phosphatase inhibitor mixture (catalog no. 78442, Thermo Fisher Scientific), 20 μM MG132, 20 μM ALLN (MilliporeSigma) using twenty-five strokes in a glass-glass Dounce homogenizer, followed by centrifugation at 1000 × g, 4°C for 5 min. The supernatant was further centrifuged at 10,000 × g, 4°C for 15 min. The supernatant was concentrated using Amicon™ Ultra-0.5 Centrifugal Filter Units (UFC500324, MilliporeSigma) and immediately stored at −80°C as the cytosolic fraction. The pellet was resuspended in RIPA lysis buffer (sc-24948, Santa Cruz Biotechnology) containing Halt protease and phosphatase inhibitor mixture, 20 μM MG132, 20 μM ALLN, 1% SDS, and 1% N-Lauryl sarcosine sodium salt as the mitochondrial fraction. The protein concentration of both fractions was determined using Pierce BCA Protein Assay Kit (ThermoFisher Scientific).

### RNA Sequencing and Transcriptomic Analysis

Total RNA was isolated from heart tissues using the Direct-zol RNA MiniPrep Plus kit (R2073, Zymo Research) following manufacturer’s instructions. For post-MI analysis, RNA was extracted from distinct cardiac regions (infarct, remote, and boundary zones) following LAD artery ligation. For diet studies, RNA was isolated from whole hearts of mice fed either standard chow or high-fat diet (HFD) for 6 months.

RNA sequencing was performed by two independent service providers. Post-MI heart tissue samples were submitted to GENEWIZ (Azenta Life Sciences) for library preparation and sequencing. Raw RNA-seq reads were first subjected to quality control and adapter trimming using Trim Galore (v0.6.5). Transcript quantification was performed using Kallisto (v0.46) with mm10 reference transcriptome. The transcript-level abundance estimates were summarized to gene-level counts using the tximport package in R. Differential expression analysis was conducted using the edgeR package (v3.34.1).

For diet studies, total RNA from chow diet and HFD groups from WT and TKO mice was processed. For each experimental group, heart tissues from two mice (50 mg/mouse) were pulverized and pooled, with three pooled biological replicates per condition. These samples were submitted to NovaGen Biotech Labs, Inc. (Great Neck, New York) for library preparation and RNA sequencing on an Illumina NovaSeq X Plus platform. Raw RNA-seq reads were processed using Trim Galore (v0.6.7) to remove adapter sequences and low-quality bases. Transcript-level quantification was performed using Kallisto (v0.46.2) with mm10 reference transcriptome and transcript abundance estimates were imported into R using the tximport package to generate gene-level count matrices. Differential expression analysis was conducted using DESeq2 (v1.32.0).

For genome browser visualization, the trimmed reads from diet studies dataset were aligned to the reference genome using HISAT2 (v2.2.1) with default parameters. Aligned reads were sorted and indexed using SAMtools and normalized coverage tracks (bigWig files) were generated using bamCoverage from deepTools (v3.5.1), using counts per million mapped reads (CPM) normalization. The tracks were visualized using the Integrative Genomics Viewer (IGV).

### Whole Genome Bisulfite Sequencing (WGBS) and DNA Methylation Analysis

High molecular weight genomic DNA was isolated from heart and testis tissues using the Monarch HMW DNA Extraction Kit (New England BioLabs, #T3060L) following manufacturer’s instructions. WGBS was performed by two independent service providers using distinct sample sets.

In the first analysis, purified WT and TKO cardiac genomic DNA samples were submitted to Azenta Life Sciences for whole genome bisulfite sequencing. The NEBNext Enzymatic Methyl-seq Kit was used for sample preparation following Azenta’s established protocols. Raw reads were first trimmed using Trim Galore (v0.6.7), with options “--adapter AGATCGGAAGAGCACACGTCTGAACTCCAGTCA --adapter2 AGATCGGAAGAGCGTCGTGTAGGGAAAGAGTGT --length 15 --clip_r1 8 --clip_r2 8 -- three_prime_clip_r1 8 --three_prime_clip_r2 8”. The trimmed reads were then aligned to bisulfite-converted mm10 reference genome using Bismark (v0.23.1) with Bowtie2 (v2.4.5) as the underlying alignment algorithm. Following alignment, duplicates were removed using deduplicate_bismark. Methylation calls were then extracted using bismark_methylation_extractor, which generated BedGraph files reporting methylation levels at single-base resolution. These deduplicated BedGraph files were converted to TDF (Tiled Data Format) files using IGVtools (v2.7.2) for visualization of DNA methylation patterns in the Integrative Genomics Viewer (IGV).

In the second analysis, high molecular weight genomic DNA was isolated from both heart and testis tissues (n=2 for each tissue type per genotype) from WT and TKO mice. These samples were submitted to Novogene for whole genome bisulfite sequencing, library preparation, and sequencing using their established protocols. Raw reads were trimmed using Trim Galore (v0.6.7), with options “--adapter AGATCGGAAGAGCGTCGTGTAGGGAAAGAGTGTAGATCTCGGTGGTCGCCGTATC ATT --adapter2 GATCGGAAGAGCACACGTCTGAACTCCAGTCACGGATGACTATCTCGTATGCCGT CTTCTGCTTG --length 15 --clip_r1 10 --clip_r2 15 --three_prime_clip_r1 10 -- three_prime_clip_r2 10”. Following trimming, the same bioinformatic pipeline was applied as described above, including alignment with Bismark, duplicate removal, methylation call extraction, and visualization preparation for IGV analysis.

### CRISPR/Cas9-Mediated Deletion of Mouse Parkin Intron 10 Differentially Methylated Region

The identified differentially methylated region (DMR) within intron 10 of the mouse Parkin (Prkn) gene was targeted for deletion. Guide RNA sequences targeting both ends of the DMR were selected using CRISPR Target Track Setting (UCSC Genome Browser), with the left and right gRNA sequences corresponding to chromosomal positions chr17:12006677-12006699 and chr17:12020834-12020856, respectively. A dual gRNA Mammalian CRISPR vector system was designed and synthesized by VectorBuilder Inc. (Chicago, IL), including both the Prkn intron 10 DMR targeting construct (pRP[2CRISPR]-EGFP/Puro-hCas9-U6>{Prkn Chr17:12006677-12006699}-U6>{Prkn Chr:12010834-12020856}) and a scrambled guide RNA control vector (pRP[CRISPR]-EGFP/Puro-hCas9-U6>Scramble_gRNA1). Sequence verification was performed by the manufacturer using Sanger sequencing. The targeting plasmid expresses Cas9 nuclease with two guide RNAs designed to delete a 14,180 bp region within intron 10 of the Parkin gene. VectorBuilder prepared the plasmids by maxiprep purification, yielding >1 μg/μl concentration (300 μl total volume).

Both VectorBuilder-produced plasmids contained EGFP and puromycin resistance markers for cellular selection and transfection tracking. 3T3-L1 fibroblast cells or C2C12 myoblast cells were seeded in 10 cm plates at approximately 50% confluency and maintained in antibiotics-free 10% FBS DMEM media. Cells were transfected with 15 μg of plasmid DNA diluted in 0.75 ml Opti-MEM I (1X, Cat# 31985062, ThermoFisher Scientific) using 60 μl EndoFectin Max Transfection Reagent (GeneCopoeia), which was also diluted in 0.75 ml Opti-MEM, following the manufacturer’s protocol. Puromycin selection (Cat. Code ant-pr-1, InvivoGen) was initiated 24 hours post-transfection and maintained for 72 hours. The selected cells were subsequently maintained in 10% FBS DMEM medium.

Genomic DNA was isolated using the GeneJet Genomic DNA Purification Kit (Thermofisher Scientific, K0721). PCR validation of the deletion was performed using three primers, which differentiate wild-type and mutant DNAs, and Go Taq™ Master Mixes 2X (Promega, M7123).

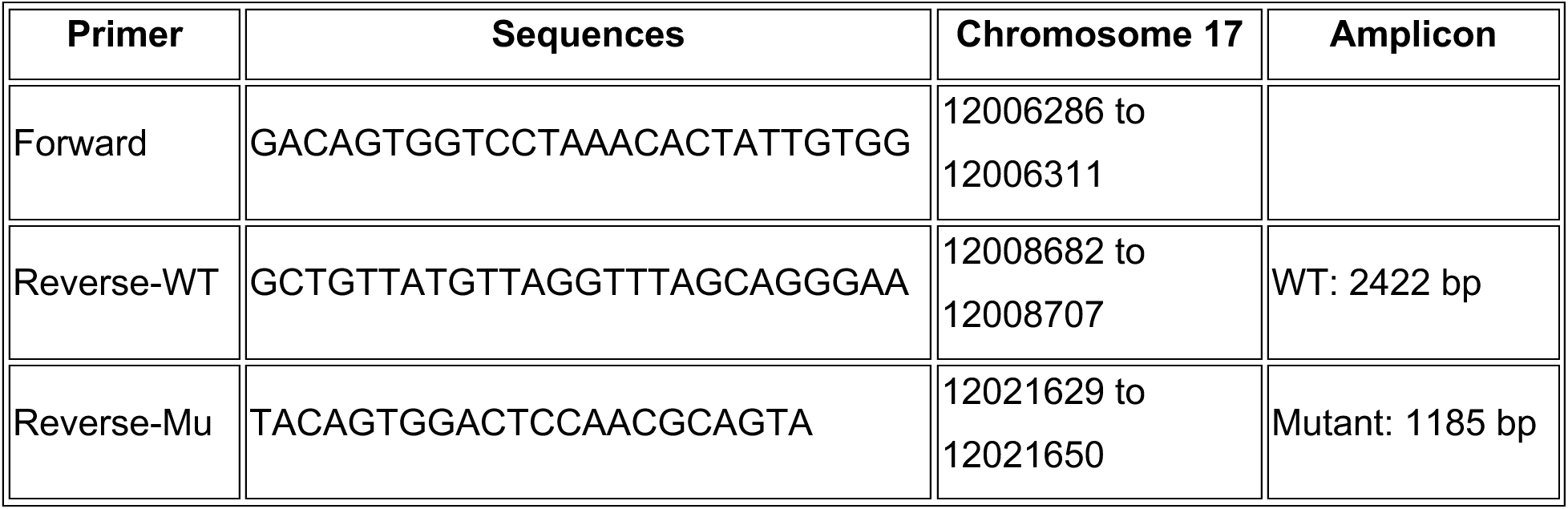

Cycling conditions for amplifying longer PCR products were used (Bench Guide, PCR, Amplification of long PCR products, Qiagen):

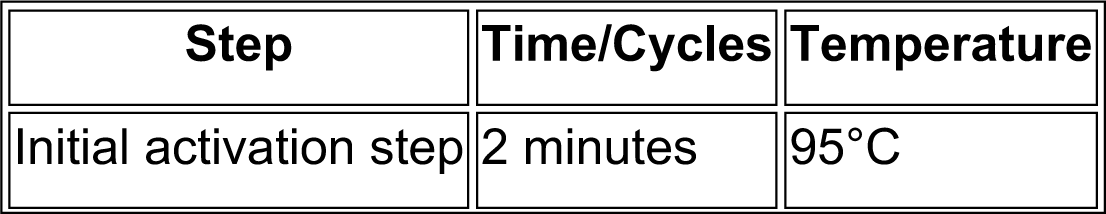
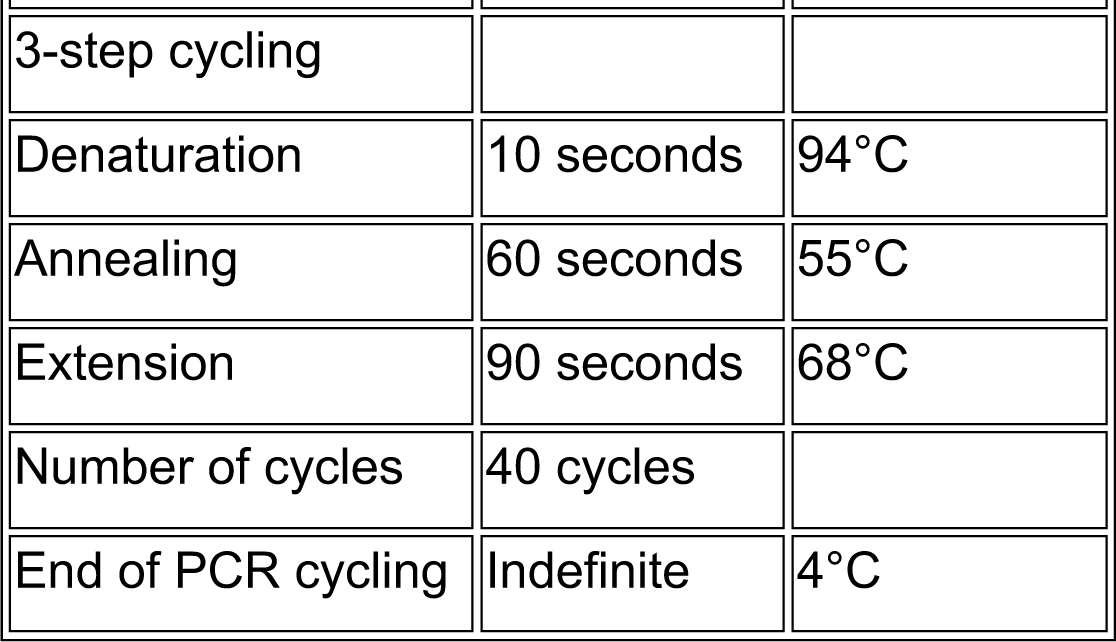

| Step | Time/Cycles | Temperature |
| --- | --- | --- |
| Initial activation step | 2 minutes | 95°C |
| 3-step cycling |  |  |
| Denaturation | 10 seconds | 94°C |
| Annealing | 60 seconds | 55°C |
| Extension | 90 seconds | 68°C |
| Number of cycles | 40 cycles |  |
| End of PCR cycling | Indefinite | 4°C |

Successful deletion yielded a 1,185 bp product compared to the 2,422 bp wild-type product when resolved on 1.0% agarose gel made in 0.5X TBE buffer. To assess the impact of DMR deletion on Parkin gene expression, RNA was extracted using Quick-RNA Miniprep (Zymo Research Inc., R1055) for Prkn mRNA quantification by qPCR. Total protein was isolated using the Total Protein Extraction Kit for Animal Cultured Cells and Tissues (Invent Biotechnologies Inc., SD-001/SN-002), followed by Western blot analysis of Parkin protein expression as described previously.

### AAV9 Vector Construction and Viral Production

Two distinct AAV9 vector systems with cardiac-specific expression were outsourced for production:

**1) AAV9-cTnT-Tigar vectors (Charles River Laboratories):** Custom AAV9 vectors expressing wild-type Tigar (NM_177003) and a phosphatase-deficient Tigar mutant were designed and manufactured by Charles River Laboratories for cardiac-specific gene delivery. The coding sequences (895 bp) were subcloned into a pAV9-cTnT-GFP backbone containing an IRES-GFP reporter. For the phosphatase-deficient mutant, three critical catalytic residues (His11, Glu102, and His198) were simultaneously substituted with alanine residues, thereby abolishing phosphatase activity while maintaining protein expression. Charles River verified the resulting constructs (pAV9-cTnT-NM_177003-IRES-GFP and pAV9-cTnT-NM_177003mutant-IRES-GFP) by full-length sequencing and performed viral packaging, yielding high-titer preparations (>1×10¹³ genomic copies/ml) delivered as 4 × 250 μl aliquots per construct. Control AAV9 viral particles containing only GFP (pAV9-cTnT-GFP, Cat# CV17196-AV9) were purchased from Charles River.
**2) AAV9-cTnT-Tigar-shRNA vectors (VectorBuilder):** For Tigar knockdown studies, an shRNA construct (pAAV[3miR30]-cTnT>EGFP:{mTigar[shRNA#1,2,3]}:WPRE) was designed and generated by VectorBuilder Inc. A corresponding control vector with a scrambled shRNA sequence (pAAV[miR30]-cTnT>EGFP:Scramble[miR30-shRNA#1]:WPRE) was also constructed by VectorBuilder. The company packaged both vectors into AAV9 particles through medium-scale production followed by ultra-purification, yielding titers exceeding 3.65×10¹³ genomic copies/ml, delivered as 10 × 50 µl aliquots per vector.

### In Vivo Delivery

For AAV9 administration, the commercially produced viral preparations were diluted to 1.2×10¹⁰ viral particles per microliter in sterile physiological saline. Mice were anesthetized with isoflurane, and 50 µl of the diluted viral suspension (total: 6×10¹¹ viral particles per mouse) was administered via retro-orbital injection. Animals were monitored post-injection for adverse events, and cardiac transgene expression was assessed four weeks post-administration.

### Quantification and statistical analyses

Prism (10) GraphPad Software was used for data processing, analyses, and graph production in the experiments. The number of independent experimental replications and the average with standard deviation are provided in the figure legends. Unpaired two-tailed t-tests or non-parametric tests (Mann Whitney, Kruskal Wallis) were used for statistical analyses. Statistical analyses were made at significance levels as follows: ns, not statistically significant; *p<0.05; **p<0.01; ***p<0.001, and ****p<0.0001.

## Supplemental Figures and Figure Legends

**Figure S1.**
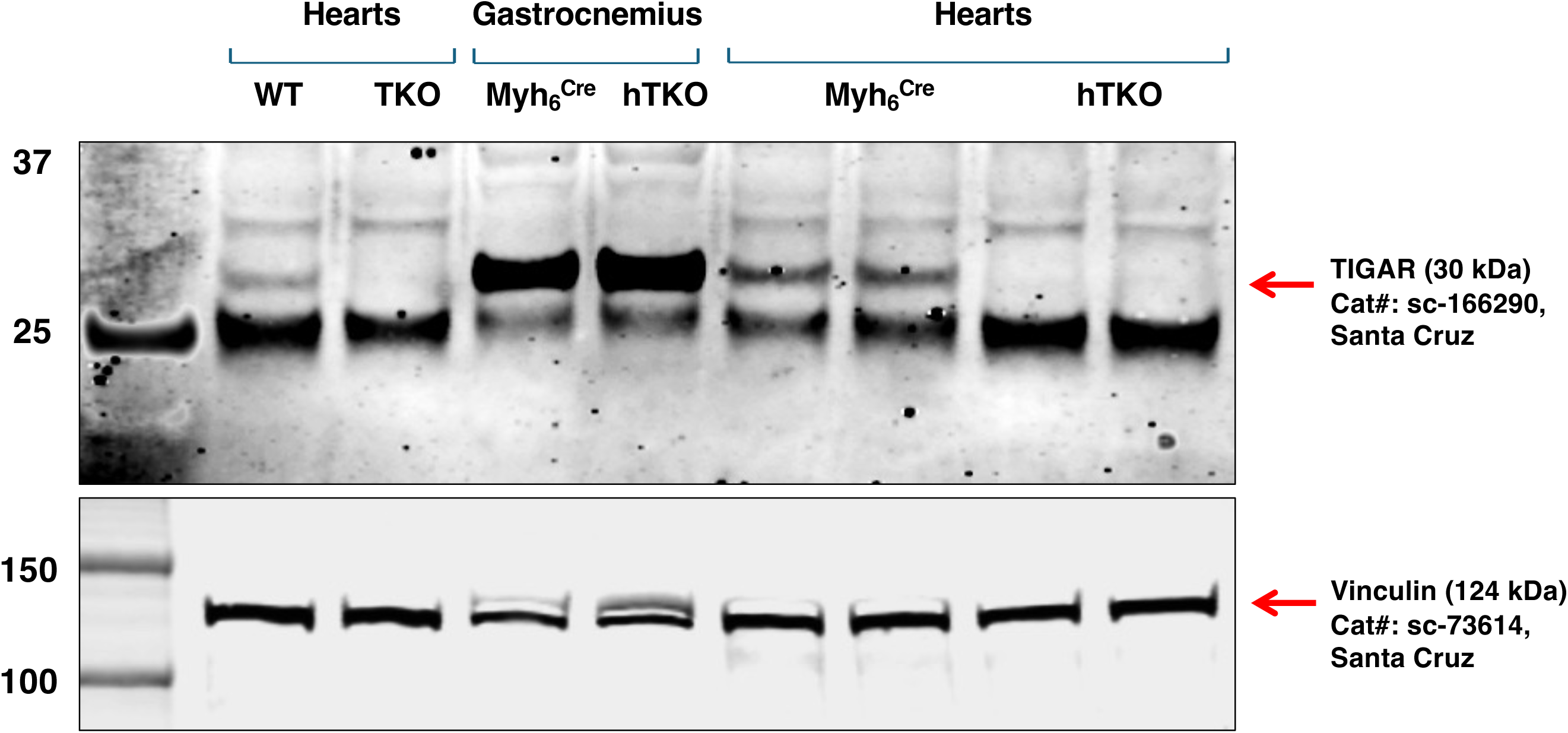
Confirmation of TIGAR knockout in heart tissues. Western blot analysis demonstrating absence of TIGAR protein (30 kDa) in TKO and hTKO hearts compared with wild-type (WT) and Myh6^Cre^ control hearts. TIGAR protein expression was comparable in gastrocnemius muscle between Myh6^Cre^ and hTKO mice, confirming cardiac-specific nature of TIGAR deletion in the hTKO model. Vinculin (124 kDa) served as loading control. **Antibodies:** TIGAR (Cat# sc-166290, Santa Cruz Biotechnology); Vinculin (Cat# sc-73614, Santa Cruz Biotechnology). TKO and hTKO samples served as negative controls for TIGAR antibody specificity.

**Figure S2.**
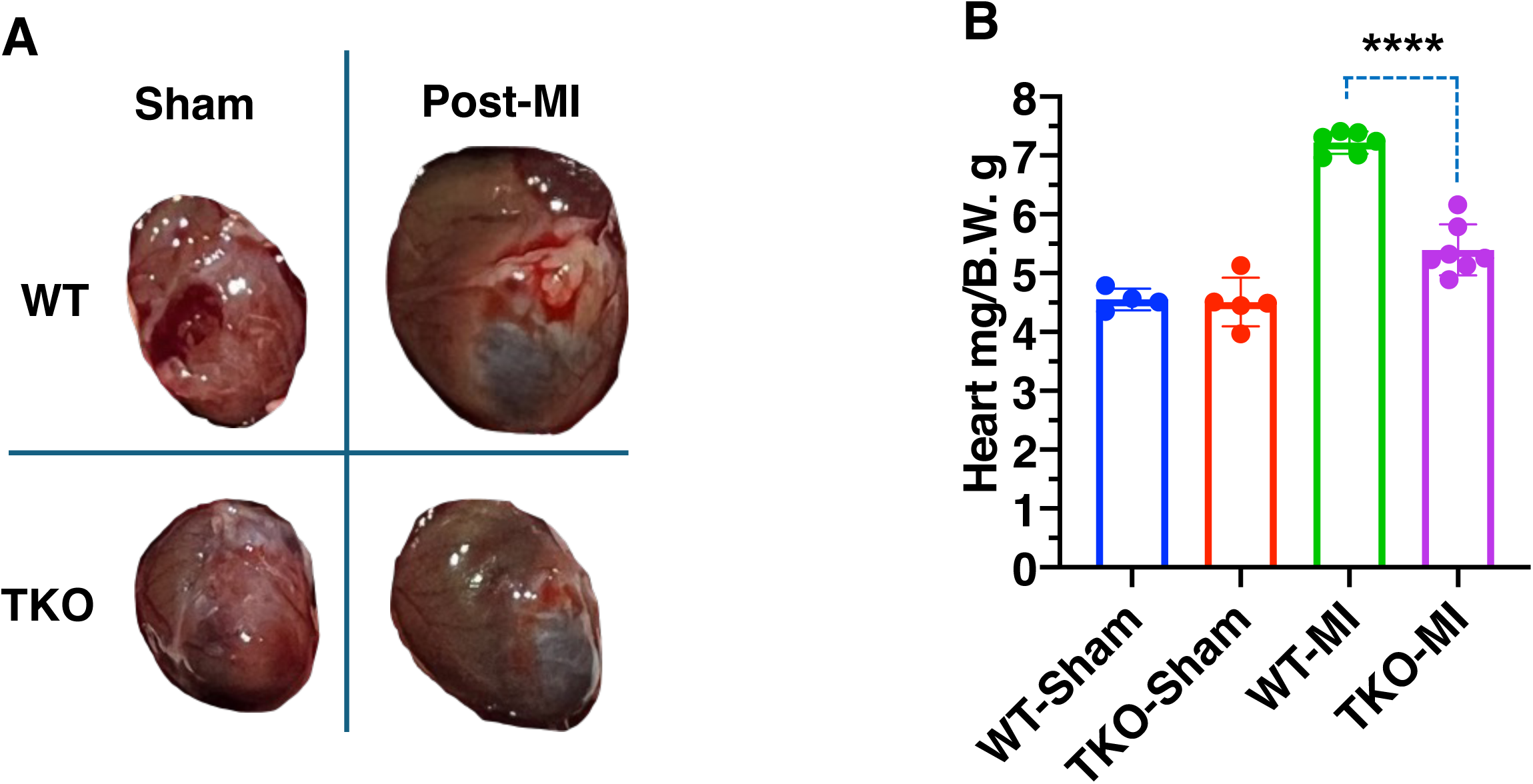
TKO mice demonstrate reduced cardiac hypertrophy following myocardial infarction. A, Representative heart images from WT and TKO mice under sham operation and post-myocardial infarction (post-MI) conditions. **B,** Heart weight-to-body weight ratios (mg/g) in experimental groups. WT mice exhibited significant cardiac hypertrophy post-MI, which was attenuated in TKO mice. Data are presented as mean±SD (n=6-7 per group). ****P<0.0001 by two-way ANOVA with Tukey’s post-hoc test.

**Figure S3.**
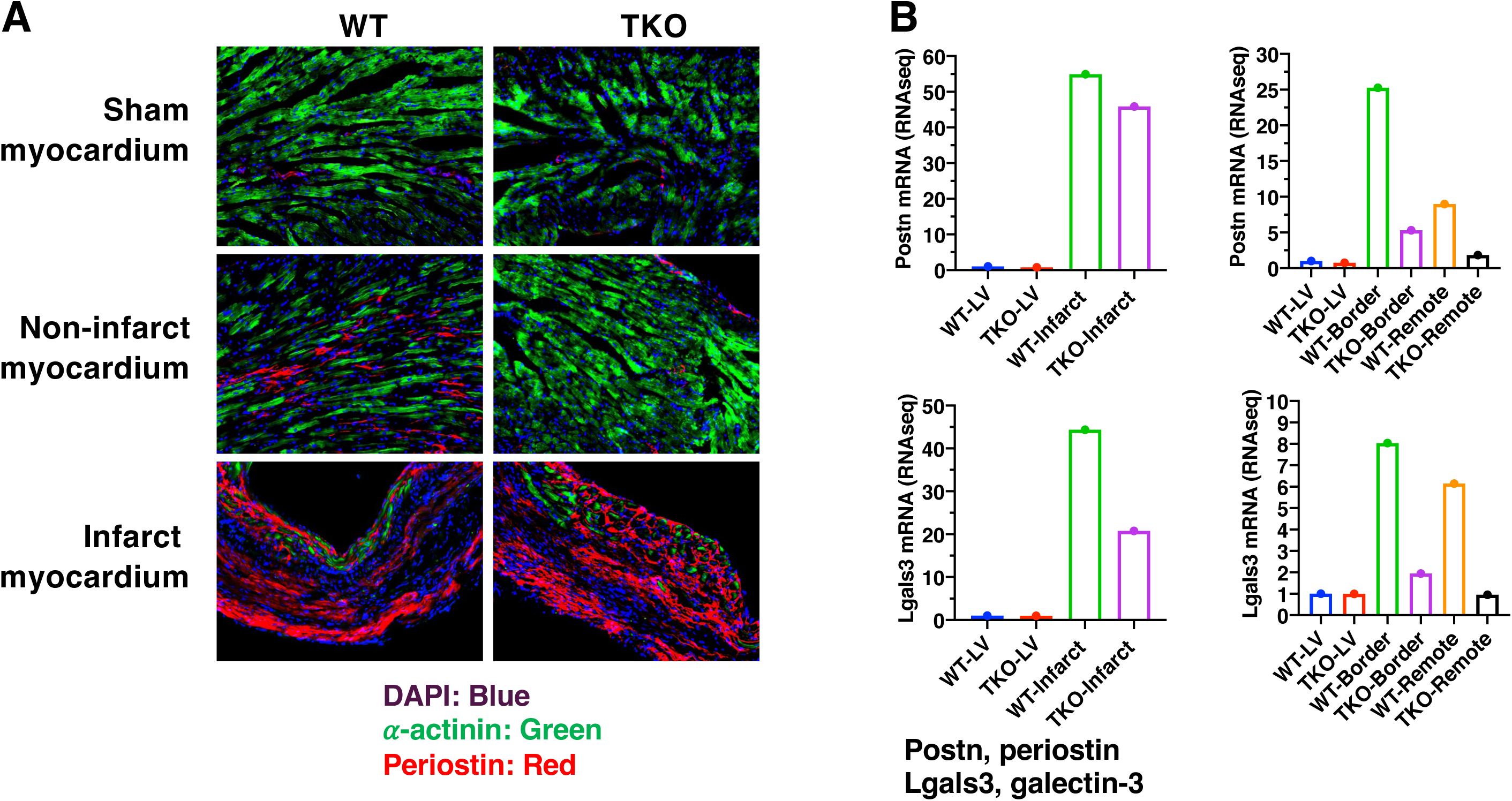
Restricted fibrosis development in TKO mice 4 weeks post-MI. A, Representative immunofluorescence images of heart sections from WT and TKO mice showing sham myocardium, non-infarct myocardium, and infarct myocardium. Sections were stained for DAPI (blue, nuclei), α-actinin (green, cardiomyocytes), and periostin (red, fibrosis marker). **B,** RNA sequencing analysis of fibrosis marker genes *Postn* (periostin) and *Lgals3* (galectin-3) in left ventricle (LV), infarct zone, border zone, and remote zone of WT and TKO hearts. TKO hearts demonstrated reduced expression of fibrosis markers in non-infarct (remote) myocardium compared with WT hearts.

**Figure S4.**
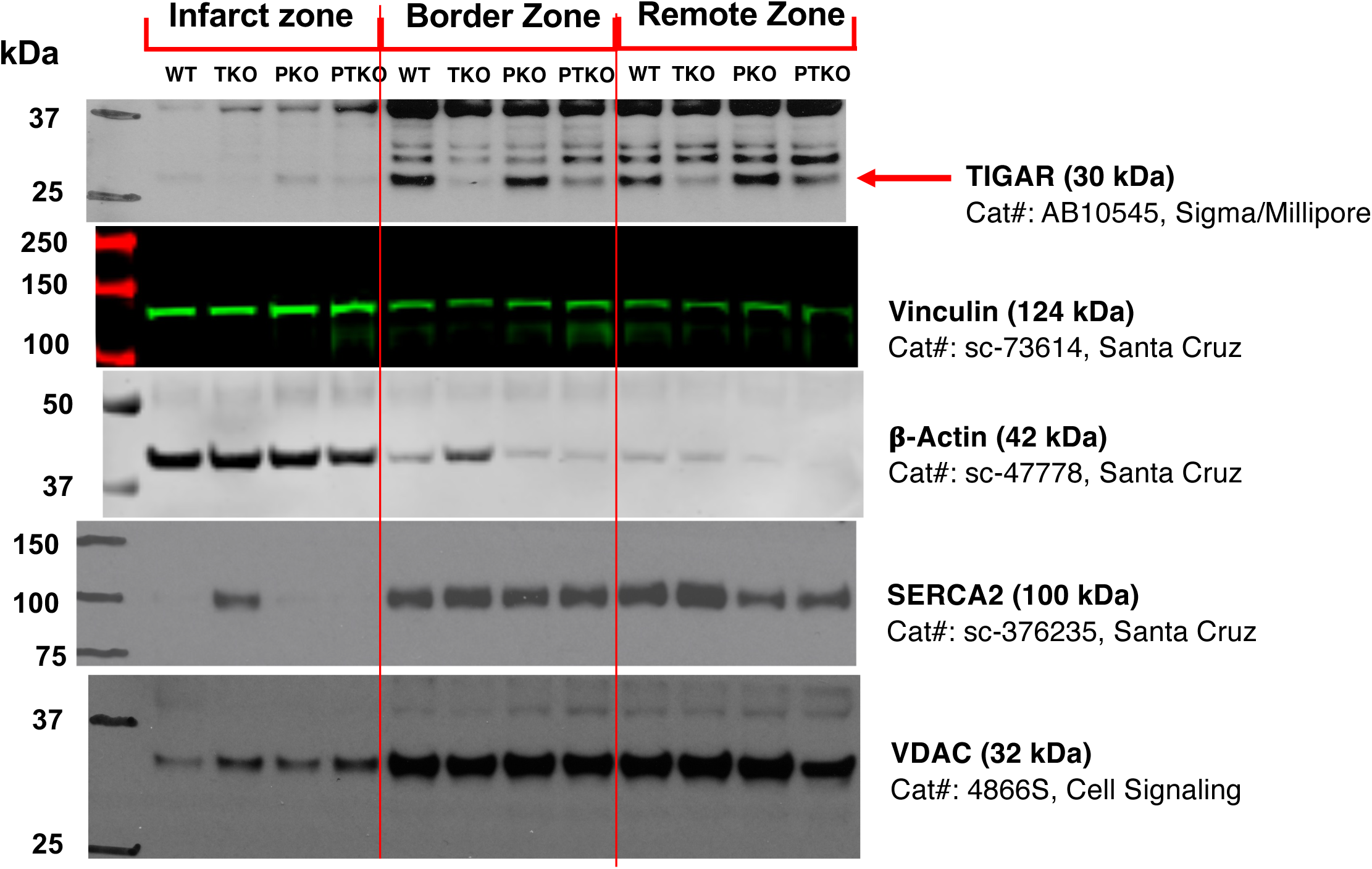
SERCA2 protein preservation in TKO heart infarct zone post-MI. Western blot analysis of protein expression in infarct zone, border zone, and remote zone from WT, TKO, Parkin knockout (PKO), and Parkin/TIGAR double knockout (PTKO) hearts post-MI. SERCA2 (100 kDa) protein was preserved in infarct zone of TKO hearts but depleted in WT, PKO, and PTKO hearts. **Tissue preparation:** Hearts were homogenized in RIPA lysis buffer (sc-24948A) containing protease/phosphatase inhibitors (Halt™), 20 μM MG-132, and 20 μM ALLN; 30 μg total protein loaded per lane. **Antibodies:** TIGAR (Cat# AB10545, Sigma/Millipore); Vinculin (Cat# sc-73614, Santa Cruz Biotechnology); β-Actin (Cat# sc-47778, Santa Cruz Biotechnology); SERCA2 (Cat# sc-376235, Santa Cruz Biotechnology); VDAC (Cat# 4866S, Cell Signaling Technology). TKO samples served as negative controls for TIGAR antibody specificity.

**Figure S5.**
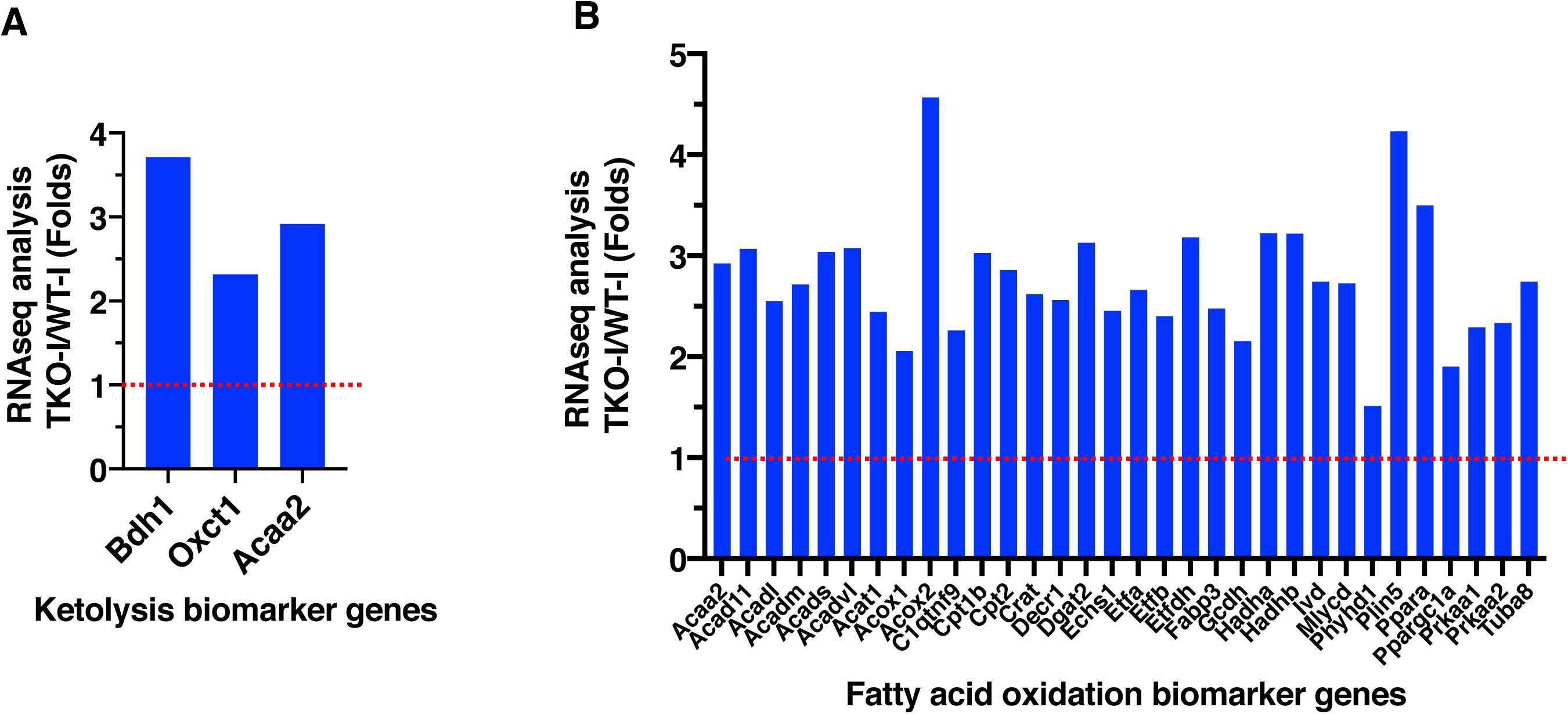
Metabolic gene expression profiles in infarct zones. RNA sequencing analysis comparing gene expression ratios (TKO-Infarct/WT-Infarct) in infarct zones. **A,** Ketolysis biomarker genes (*Bdh1*, *Oxct1*, *Acaa2*). **B,** Fatty acid oxidation biomarker genes. Results demonstrate relative preservation of metabolic gene expression in TKO infarct zones compared with WT infarct zones. While these metabolic genes were downregulated in infarct zones of both genotypes compared with non-infarct tissue, fold changes >1 indicate that TKO hearts maintained relatively higher expression levels, suggesting preserved metabolic capacity following myocardial infarction.

**Figure S6.**
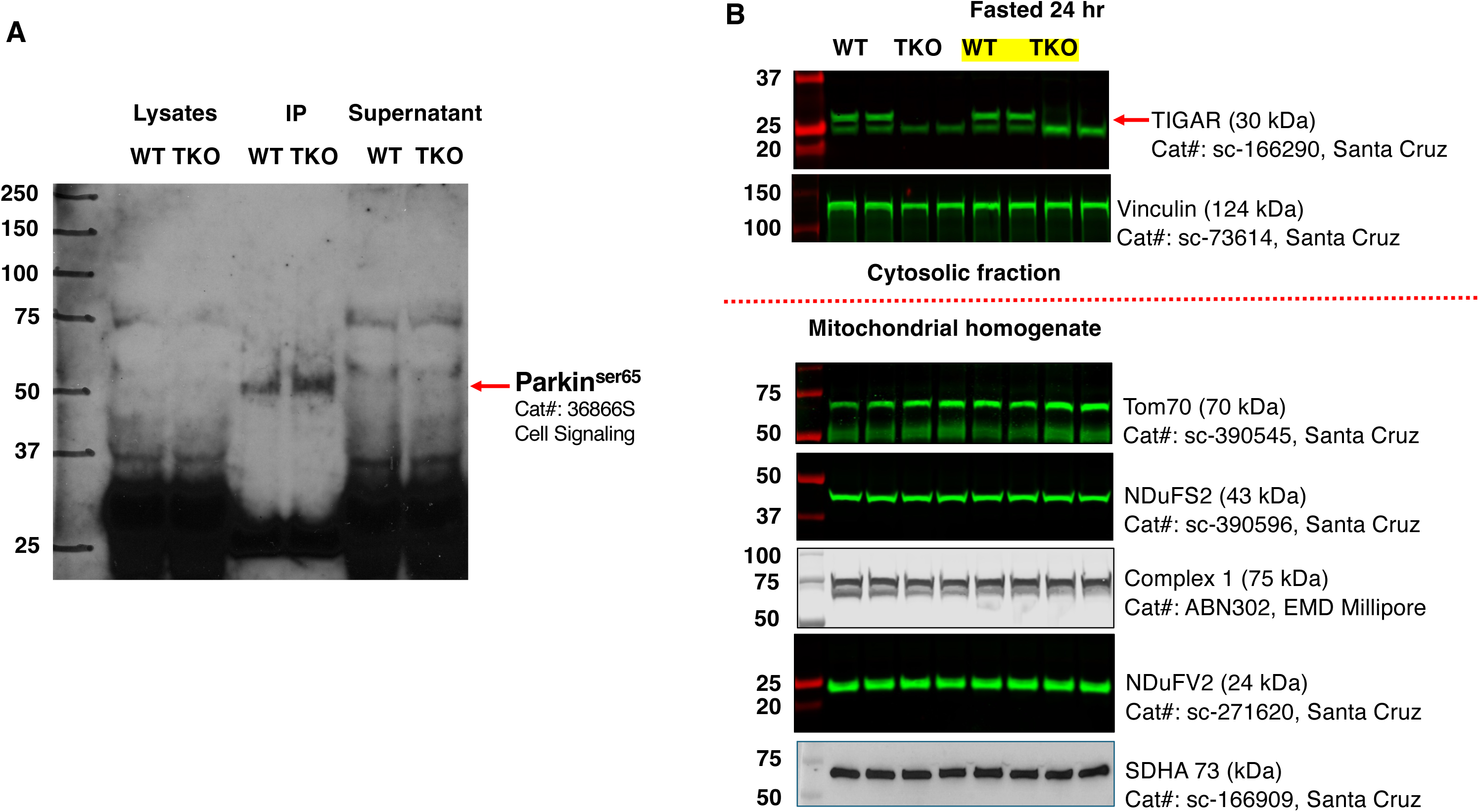
Parkin phosphorylation and mitochondrial protein analysis in TKO hearts. A, Analysis of Parkin serine 65 phosphorylation in WT and TKO hearts. Heart tissues were homogenized in RIPA lysis buffer (composition as described in Supplementary Figure 4), and 2 mg lysate was subjected to immunoprecipitation (IP). Lysate lanes show 40 μg total protein (2% input) before immunoprecipitation. IP lanes represent samples immunoprecipitated with Parkin antibody and immunoblotted with phospho-Parkin Ser65 antibody. Supernatant lanes show post-IP samples (40 μg). **B,** Western blot analysis of mitochondrial proteins in cytosolic fractions and mitochondrial homogenates from fed and 24-hour fasted WT and TKO mice. Proteins analyzed include TIGAR (30 kDa), Vinculin (124 kDa), Tom70 (70 kDa), NDUFS2 (43 kDa), Complex I (75 kDa), NDUFV2 (24 kDa), and SDHA (73 kDa). Comparable levels of mitochondrial proteins were observed between WT and TKO hearts under normal conditions. **Antibodies:** Parkin (Cat# sc-32282, Santa Cruz Biotechnology); phospho-Parkin Ser65 (Cat# 36866S, Cell Signaling Technology); TIGAR (Cat# sc-166290, Santa Cruz Biotechnology); Tom70 (Cat# sc-390545, Santa Cruz Biotechnology); NDUFS2 (Cat# sc-390596, Santa Cruz Biotechnology); Complex I (Cat# ABN302, EMD Millipore); NDUFV2 (Cat# sc-271620, Santa Cruz Biotechnology); SDHA (Cat# sc-166909, Santa Cruz Biotechnology).

## Notes

### Competing Interest Statement

The authors have declared no competing interest.

